# Improving management strategies of plant diseases using sequential sensitivity analyses

**DOI:** 10.1101/315747

**Authors:** Loup Rimbaud, Sylvie Dallot, Claude Bruchou, Sophie Thoyer, Emmanuel Jacquot, Samuel Soubeyrand, Gaël Thébaud

## Abstract

Improvement of management strategies of epidemics is often hampered by constraints on experiments at large spatiotemporal scales. A promising approach consists of modelling the biological epidemic process and human interventions, which both impact disease spread. However, few methods enable the simultaneous optimisation of the numerous parameters of sophisticated control strategies. To do so, we propose a heuristic approach (i.e., a practical improvement method approximating an optimal solution) based on sequential sensitivity analyses. In addition, we use an economic improvement criterion, based on the net present value, accounting for both the cost of the different control measures and the benefit generated by disease suppression. This work is motivated by sharka (caused by *Plum pox virus*), a vector-borne disease of prunus trees (especially apricot, peach and plum) whose management in orchards is mainly based on surveillance and tree removal. We identified the key parameters of a spatiotemporal model simulating sharka spread and control, and approximated optimal values for these parameters. The results indicate that the current French management of sharka efficiently controls the disease, but can be economically improved using alternative strategies that are identified and discussed. The general approach should help policymakers to design sustainable and cost-effective strategies for disease management.

## INTRODUCTION

Improving large-scale disease management constitutes a major challenge. Faced with the urgent need to deal with emerging epidemics, one often relies on expert opinions to design management strategies, but opinions are not necessarily based on quantitative data. Some specific control methods can be tested through field trials, but they constitute only part of a global wider management strategy, which must be assessed at large spatiotemporal scales. However, at such scales field trials are considerably limited by tractability issues, resulting in poorly generalizable results from the very few trials that might be carried out. Epidemiological models have been very helpful to overcome these obstacles and to account for the interactions between biological processes and human interventions that jointly impact disease spread (Jeger et al. 2018; Parnell et al. 2017). The key epidemiological parameters of these models are prime targets for control measures. These parameters can be identified using global sensitivity analysis, as shown for invasive plants (Coutts et al. 2011), and plant pathogens such as viruses (Chan and Jeger 1994; Holt et al. 1999; Jeger and Chan 1995; Rimbaud et al. 2018a), fungi (Papaïx et al. 2014; Xu and Ridout 1998), and bacteria (Breukers et al. 2007). By varying input parameters of a simulation model within their respective variation ranges (corresponding to their natural variation or to the uncertainty on their estimates), global sensitivity analysis allows the computation of ‘sensitivity indices’ quantifying the influence of the variability of each input parameter on a given output variable (Saltelli et al. 2008). The definition of parameter variation ranges depends on the objectives of the modeller (e.g., identifying key drivers of a biological process, identifying parameters which deserve to be better estimated), and are informed by quantitative data or expert opinions.

Some epidemiological models are used to assess the potential of different control actions in various epidemic scenarios. In this context, studies dealing with the management of perennial plant diseases have addressed management strategies based on roguing and possibly replanting (Cunniffe et al. 2014, 2015, 2016; Filipe et al. 2012; Hyatt-Twynam et al. 2017; Ndeffo Mbah and Gilligan 2010; Parnell et al. 2009, 2010; Sisterson and Stenger 2013), planting with different densities (Chan and Jeger 1994; Cunniffe et al. 2014, 2015; Jeger and Chan 1995), planting outside of contaminated areas (Chan and Jeger 1994; Filipe et al. 2012; Jeger and Chan 1995), or spraying with insecticides (Filipe et al. 2012). These studies optimised one or two control parameters under various epidemiological scenarios, but the other control parameters remained fixed at their reference value. Because control parameters generally interact each other, it is crucial to develop an alternative approach that jointly explores numerous combinations of control parameters to identify promising combinations of parameter values.

Our work is motivated by the management of sharka, the most damaging disease of prunus trees (Cambra et al. 2006). Its causal agent, *Plum pox virus* (PPV, genus *Potyvirus*), is naturally transmitted by more than 20 aphid species in a non-persistent manner (Labonne et al. 1995) and has spread worldwide due to human shipping and planting of infected material (Cambra et al. 2006). Faced with the threat posed by sharka, various management strategies have been adopted in different countries (Rimbaud et al. 2015). In countries targeting disease limitation (e.g., France) or eradication (e.g., United States), a key element of the management strategy is the appropriate surveillance of nurseries and orchards, followed by tree removals. Surveillance methods rely on leaf sampling followed by serological or molecular diagnostic tests or, alternatively, visual inspection of prunus trees to detect sharka symptoms (mostly on leaves and fruits). When infected trees are detected, they are culled to remove sources of inoculum. Because of its long incubation period, PPV infection may remain undetected for several months or years after inoculation, regardless of the surveillance method (Quiot et al. 1995; Sutic 1971). Combined with evidence of short-distance transmission of PPV (Dallot et al. 2003; Gottwald et al. 2013; Pleydell et al. 2018), this delay explains why non-symptomatic trees surrounding detected trees may need to be removed as well. In France, sharka management is compulsory and a national decree specifies the control actions mentioned above (JORF 2011). In particular, this decree describes a procedure defining the frequency of visual surveillance of orchards depending on their distance to the nearest detected infection, the removal of whole orchards if their annual contamination rate exceeds a given threshold, and the conditions for replanting (Fig. 4A). This strategy is based on expert opinions and not on quantitative data or a formal demonstration of its efficiency. In addition, it is expensive (and complex) to implement. It is thus crucial to conduct a formal cost-benefit analysis and identify improved management rules.

The three objectives of this work are: i) the development of a heuristic approach (i.e., a practical method, not guaranteed to be optimal, but instead sufficient for reaching a goal) to improve management strategies of epidemics using sequential sensitivity analyses; ii) the design of an economic criterion which accounts for the balance between the costs induced by control actions and the benefits of reducing epidemic damage; and iii) the identification of economically improved alternatives to the current French strategy to manage sharka. To do so, we use a spatiotemporal model initially developed in previous studies (Pleydell et al. 2018; Rimbaud et al. 2018a). This model simulates the turnover of peach orchards in a real landscape, recurrent PPV introductions at planting and PPV spread by aphid vectors; a previous article presented an in-depth analysis of the model sensitivity to the included epidemiological parameters (Rimbaud et al. 2018a). In the present work, we implemented in this model an explicit management strategy, which is flexible enough to encompass sharka management in several countries including France. Consequently, the identified strategies should be of interest to the countries targeting area-wide management of sharka and more generally perennial plant diseases.

## MATERIALS AND METHODS

### Model description

#### Model overview

We use a stochastic, spatially-explicit, SEIR (susceptible-exposed-infectious-removed) model developed to simulate sharka epidemics in a real cultivated landscape (Pleydell et al. 2018; Rimbaud et al. 2018a). This landscape, typical of a peach-growing area in southeastern France, comprises 553 patches (i.e., land parcels, which are not necessarily contiguous), representing a total area of 524 ha (average patch area: 0.95 ha; see S1 map in Rimbaud et al. 2018a). Each patch is cultivated with a succession of peach orchards (i.e., a peach crop planted at a given date, with an average density of approximately 720 trees.ha^-1^). The model is patch-based, with a discrete time step of 1 week. Each host is in one of five health states (Fig. 1A). The epidemic process and the transitions between states were described previously (Pleydell et al. 2018; Rimbaud et al. 2018a). Briefly, at the beginning of the simulation, PPV is introduced for the first time (via orchard planting) in a single patch identified using its connectivity with other patches, *q_κ_*. We define the connectivity of a patch as the mean number of infectious aphids that would leave this patch if all trees in this patch were infectious, and land in the other patches of the landscape. Thus, *q_κ_* depends on the patch area and proximity to other patches and relates to its potential to initiate an epidemic. Further introductions can occur at every orchard planting with probability *Φ*. At each introduction, the prevalence (i.e., the proportion of infected trees in the orchard at the time of planting) *τ* is drawn from a mixture distribution favouring high or low prevalences depending on the relative probability of massive introduction, *p_MI_* Once introduced in orchards, the pathogen is spread by aphid vectors at a distance that depends on their dispersal kernel, parameterised by the mean (*W_exp_*) and the variance (*W_var_*) of a dispersal variable (see Rimbaud et al. 2018a for details). Healthy trees (susceptible state S, Fig. 1A) become infected (exposed state E) when successfully inoculated by infectious aphid vectors (the probability of infection depends on the aphid transmission coefficient *β*). After a latent period (drawn from a Gamma distribution with mean duration *θ_exp_* and variance *θ_var_* and truncated in such way that the latent period cannot end within the growing season in which the host is infected), infected trees become infectious and symptomatic (hidden state H, i.e., these trees are not yet detected), from where they may be detected (state D, with probability *ρ*). Once detected, hosts are culled (removed state R) after a mean delay *δ*, and at the latest by the end of the year (using a truncated Geometric distribution). It is assumed that infectious (i.e., symptomatic) trees are no longer productive, due to diseased-induced fruit loss or ban on fruit sales. However, the disease is not supposed to affect host lifespan (no available data report any increase in prunus mortality due to PPV), thus hosts can reach state R only if removed. Due to orchard turnover (independent of the disease, simulated by drawing orchard duration from a Poisson distribution with mean *ψ*=15 years), whole orchards can be replaced by trees in state S (or possibly H). Table 1 summarises all the model parameters.

**Fig. 1.**
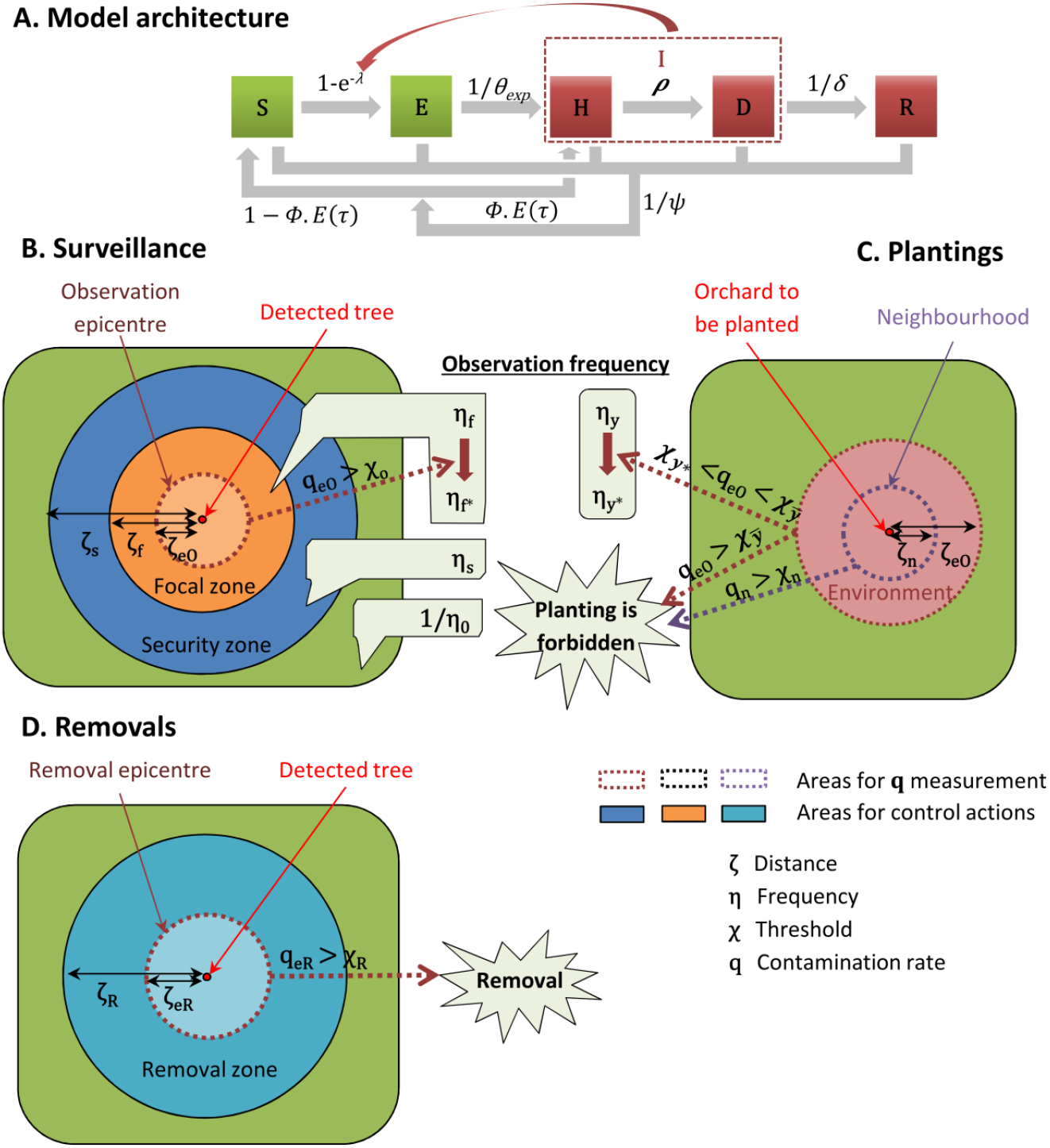
Schematic representation of the spatiotemporal stochastic model simulating sharka spread and management. (A) Flow diagram of the SEHDR architecture. S: susceptible (i.e., healthy); E: exposed (i.e., infected but neither infectious nor diseased); I: infectious and symptomatic; H: hidden (i.e., infectious but not yet detected); D: detected (and infectious); R: removed. Productive and non-productive (diseased) compartments are represented in green and red, respectively. (B) Surveillance of orchards, depending on their location relative to previously infected trees and on the contamination rate of the observation epicentre. (C) Planting restrictions and surveillance for young orchards, depending on the contamination rates of the environment and neighbourhood. (D) Removal of orchards, depending on the contamination rate of the removal epicentre. In the French sharka management strategy, the removal zone and its epicentre correspond to the orchard only (i.e., *ζ_eR_* = *ζ_R_* = 0; see also Fig. 4A). All model parameters are defined in the text and listed in Table 1.

**TABLE 1.** Model parameters: description, reference values for sharka epidemics in French peach orchards, and variation ranges in the sensitivity analyses. The first sensitivity analysis (step 1) targeted 23 control parameters and 6 epidemiological parameters to assess their relative influence on model outputs. In the second step, only the 17 most influential control parameters (bounds in bold) were kept and improved. In the third step, 6 unnecessary parameters were removed, 1 parameter was fixed, and the remaining 10 parameters (in bold) were further improved. In the second and third steps, the epidemiological parameters were not targeted in the sensitivity analysis, but for each simulated epidemic they were drawn from a uniform distribution within the variation range used in the first step.

| Parameter and description |  | Reference value | Steps 1 and 2 |  | Step 3 |  |
| --- | --- | --- | --- | --- | --- | --- |
|  |  |  | Min | Max | Min | Max |
| Temporal and landscape parameters |  |  |  |  |  |  |
| $a_m$ | First year of management | 6 | - | - | - | - |
| $a_f$ | Last year of simulation | 35 | - | - | - | - |
| $\psi$ | Expected orchard duration (years) | 15 <sup>a</sup> | - | - | - | - |
| Epidemiological parameters |  |  |  |  |  |  |
| $q_\kappa$ | Quantile of the connectivity of the patch of first introduction | 0.50 | 0.00 <sup>b</sup> | 1.00 <sup>b</sup> | - | - |
| $\phi$ | Probability of introduction at planting | 0.007 <sup>a</sup> | 0.0046 <sup>c</sup> | 0.0107 <sup>c</sup> | - | - |
| $p_{MI}$ | Relative probability of massive introduction | 0 <sup>a</sup> | 0.00 <sup>a</sup> | 0.10 <sup>a</sup> | - | - |
| $W_{exp}$ | Expected value of the dispersal variable | 0.486 <sup>c</sup> | 0.469 <sup>c</sup> | 0.504 <sup>c</sup> | - | - |
| $W_{var}$ | Variance of the dispersal variable | 0.0434 <sup>c</sup> | - | - | - | - |
| $\beta$ | Transmission coefficient | 1.32 <sup>c</sup> | 1.25 <sup>c</sup> | 1.39 <sup>c</sup> | - | - |
| $\theta_{exp}$ | Expected duration of the latent period (years) | 1.92 <sup>c</sup> | 1.71 <sup>c</sup> | 2.14 <sup>c</sup> | - | - |
| $\theta_{var}$ | Variance of the latent period duration (years<sup>2</sup>) | 0.44 <sup>c</sup> | - | - | - | - |
| Surveillance parameters |  |  |  |  |  |  |
| $\rho$ | Probability of detection of a symptomatic tree | 0.66 <sup>c</sup> | 0 <sup>b</sup> | 0.66 <sup>h</sup> | 0.2 | 0.66 |
| $\gamma_o$ | Duration of surveillance zones (years) | 3 <sup>d</sup> | 0 <sup>b</sup> | 10 <sup>i</sup> | 0 | 10 |
| $\gamma_y$ | Duration of young orchards (years) | 3 <sup>d</sup> | 0 <sup>b</sup> | 10 <sup>i</sup> | 0 | 3 |
| $\zeta_s$ | Radius of security zones (m) | 2,500 <sup>d</sup> | 0 <sup>b</sup> | 5,800 <sup>j</sup> | 0 | 1,000 |
| $r_{\zeta_f}$ | Ratio of the focal area over the security area | 0.36 <sup>d</sup> | 0 <sup>b</sup> | 1 <sup>b</sup> | - | - |
| $r_{\zeta_{eo}}$ | Ratio of the observation epicentre area over the focal area | 0.14 <sup>d</sup> | 0 <sup>b</sup> | 1 <sup>b</sup> | - | - |
| $1/\eta_o$ | Maximal period between 2 observations (years) | 6<sup>d</sup> | 1<sup>b</sup> | 15<sup>k</sup> | 1 | 15 |
| $\eta_s$ | Surveillance frequency in security zones (year<sup>-1</sup>) | 1<sup>d</sup> | 0<sup>b</sup> | 8<sup>l</sup> | 0 | 4 |
| $\eta_f$ | Surveillance frequency in focal zones (year<sup>-1</sup>) | 2<sup>d</sup> | 0<sup>b</sup> | 8<sup>l</sup> | - | - |
| $\eta_{f*}$ | Modified surveillance frequency in focal zones (year<sup>-1</sup>) | 3<sup>d</sup> | 0<sup>b</sup> | 8<sup>l</sup> | - | - |
| $\eta_y$ | Surveillance frequency in young orchards (year<sup>-1</sup>) | 2<sup>d</sup> | 0<sup>b</sup> | 8<sup>l</sup> | 0 | 2 |
| $\eta_{y*}$ | Modified surveillance frequency in young orchards (year<sup>-1</sup>) | 3<sup>d</sup> | 0<sup>b</sup> | 8<sup>l</sup> | - | - |
| $\chi_o$ | Contamination threshold in the observation epicentre, above which surveillance frequency in the focal zone is modified | 0.02<sup>d</sup> | 0<sup>b</sup> | 1<sup>b</sup> | - | - |
| $r_{\chi_{y*}}$ | Ratio (over $\chi_{\bar{y}}$) of the contamination threshold in the environment, above which surveillance frequency in young orchards is modified | 0.50<sup>d</sup> | 0<sup>b</sup> | 1<sup>b</sup> | - | - |
| **Removal parameters** |  |  |  |  |  |  |
| $\delta$ | Mean delay before removal of a detected tree (days) | 10<sup>d</sup> | - | - | - | - |
| $Y_{R^T}$ | (Boolean) Individual trees are removed:<br>0: after a mean delay $\delta$; 1: at the end of the year | 0<sup>d</sup> | 0<sup>b,m</sup> | 1<sup>b,m</sup> | 0 | |
| $Y_R$ | (Boolean) Whole orchards are removed:<br>0: after a mean delay $\delta$; 1: at the end of the year | 1<sup>d</sup> | 0<sup>b,m</sup> | 1<sup>b,m</sup> | - | - |
| $\zeta_R$ | Radius of the removal zone (m) | 0<sup>d</sup> | 0<sup>b</sup> | 5,800<sup>j</sup> | - | - |
| $r_{\zeta_{eR}}$ | Ratio of the removal epicentre area over the removal area | 0<sup>d</sup> | 0<sup>b</sup> | 1<sup>b</sup> | - | - |
| $\chi_R$ | Contamination threshold in the removal epicentre, above which orchards inside the removal zone are removed | 0.10<sup>d</sup> | 0<sup>b</sup> | 0.34<sup>n</sup> | - | - |
| **Replanting parameters** |  |  |  |  |  |  |
| $\gamma_s$ | Delay before replanting a removed orchard (years) | 0<sup>d</sup> | 0<sup>b</sup> | 10<sup>i</sup> | 0 | 7 |
| $\zeta_n$ | Radius of the neighbourhood (m) | 200<sup>d</sup> | 0<sup>b</sup> | 5,475<sup>j</sup> | 0 | 5,475 |
| $\chi_{\bar{y}}$ | Contamination threshold in the environment around young orchards, above which the planting of orchards is forbidden | 0.02<sup>d</sup> | 0<sup>b</sup> | 1<sup>b</sup> | - | - |
| $\chi^n$ | Contamination threshold in the neighbourhood, above which the planting of orchards is forbidden | 0.05<sup>d</sup> | 0<sup>b</sup> | 1<sup>b</sup> | 0.2 | 1 |
| **Economic parameters** |  |  |  |  |  |  |
| $y$ | Relative age-dependent yield of hosts in S or E states | 0.00 until 2 years<sup>a</sup><br>0.50 at 3 years<sup>a</sup><br>0.65 at 4 years<sup>a</sup> | - | - | - | - |

|  |  | 0.85 at 5 years <sup>a</sup> |  |  |  |  |
| --- | --- | --- | --- | --- | --- | --- |
|  |  | 1.0 from 6 to 15 years <sup>a</sup> |  |  |  |  |
|  |  | 0.80 from 16 years <sup>a</sup> |  |  |  |  |
| $c_S$ | Planting cost of one orchard (€·ha<sup>-1</sup>) | 14,000 <sup>a</sup> | - | - | - | - |
| $c_O$ | Cost of one observation (€·ha<sup>-1</sup>) | 160 <sup>a,f</sup> | - | - | - | - |
| $c_R^T$ | Removal cost of one individual tree (€) | 15 <sup>a</sup> | - | - | - | - |
| $c_R$ | Removal cost of one orchard (€·ha<sup>-1</sup>) | 1,000 <sup>a</sup> | - | - | - | - |
| $c_H$ | Yearly harvest cost (€·ha<sup>-1</sup>) | 16,500 <sup>a</sup> | | | | |
| $c_F$ | Yearly fixed cost associated with prunus cultivation (€·ha<sup>-1</sup>) | 13,600 <sup>e</sup> | - | - | - | - |
| $p$ | Maximal yearly benefit generated by fruit harvest (€·ha<sup>-1</sup>) | 37,250 <sup>e</sup> | - | - | - | - |
| $\tau_a$ | Discount rate | 0.04 <sup>g</sup> | - | - | - | - |
| $\chi_{SEHD}$ | Minimum proportion of living trees, below which the orchard is not profitable and thus removed | 0.66 <sup>e</sup> | - | - | - | - |

<sup>a</sup> estimated through expert opinion (i.e., professional organisations in charge of sharka management in France).
<sup>b</sup> mathematical domain of definition.
<sup>c</sup> estimated in (Pleydell et al. 2018).
<sup>d</sup> fixed by (JORF 2011). With regard to $\zeta_R$ and $r_{\zeta_{eR}}$ , when fixed at 0, the removal zone or the removal epicentre correspond to the contaminated orchard only.
<sup>e</sup> estimated from the economic analysis of prunus cultivation (supplementary Table S1).
<sup>f</sup> with $\rho=0.66$ .
<sup>g</sup> (Quinet 2013).
<sup>h</sup> we assumed that the detection probability could not be higher because of technical limitations.
<sup>i</sup> these durations aim to detect latent infections, and 10 years is well beyond the expected duration of the latent period ( $\theta_{exp}$ ).

In this work, we complemented the existing model with an explicit disease management strategy. After an initial epidemic period (arbitrarily fixed at five years to enable pathogen establishment), a strategy based on orchard surveillance, tree removals and replanting restrictions is applied during 30 years (which is twice the mean lifespan of the hosts and consequently a reasonable duration in which to assess the efficiency of a management strategy). The reference management strategy is based on the French management of sharka in prunus orchards (JORF 2011). Nevertheless, our model allows a multitude of variations on each control measure, which makes it possible to include some aspects of sharka management in other countries (e.g., removal radius around detected trees as in the United States; Gottwald et al. 2013). The simulated management strategy is defined by 24 parameters (14 for surveillance, 6 for removals and 4 for replanting restrictions; see Fig. 1 and Table 1).

Each year, orchard surveillance, removals and planting bans depend on the location of the corresponding patch relative to previously infected trees, and the local contamination rate in the preceding year. Let *Ω* denote the set of orchards composing the area where the contamination rate is measured (i.e., all orchards cultivated in patches whose border is within a given radius of the centroid of the focal patch), 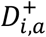 the number of trees detected in orchard *i* (*i*=1,…,*I*) during year *a* (*a*=1,…,*a_f_*), and *N_i_* the initial number of trees planted in this orchard. Then the contamination rate in this area for year *a* is:

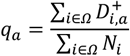

Note that this measure is neither an annual incidence (i.e., the number of detected trees divided by the number of trees at the beginning of the year) nor a prevalence, but the number of detected trees divided by the number of trees at orchard planting, which is easier to calculate directly by plant health services in the field.

#### Orchard surveillance

All orchards are surveyed visually for disease at least once every 1/*η_0_* years (hence *η_0_* is the basic surveillance frequency), and at each survey symptomatic hosts (in state H) are independently detected with probability *ρ*. Detection of a symptomatic tree triggers the definition (for the *γ_o_* following years) of nested concentric zones around the corresponding orchard (Fig. 1B). Within the focal zone, whose radius is *ζ_f_*, orchards are surveyed *η*_f_ times per year. Furthermore, an ‘observation epicentre’, which is nested within the focal zone and whose radius is *ζ_eO_* (*ζ_eO_* < *ζ_f_*), is defined to calculate the contamination rate (noted *q_eO_*). If the contamination rate of the observation epicentre exceeds a threshold *χ_o_*, the surveillance frequency in the focal zone is changed to *η_f_\**. Within the security zone, which extends from the outer boundary of the focal zone to a radius *ζ_s_* (*ζ_s_* > *ζ_f_*), orchards are surveyed *η_s_* times per year.

Additionally, young orchards are surveyed *η_y_* times per year during *γ_y_* years, but this frequency is changed to *η_y*_* if, the year before orchard planting, the contamination rate of the environment around the patch to be planted exceeds a threshold value *χ_y*_* (Fig. 1C). The environment is a zone around the patch to be planted; its radius is *ζ_eO_* (i.e., the same as the observation epicentre).

When different surveillance frequencies are assigned to a single orchard (e.g., because it is young and located in the intersection of focal and security zones defined around different contaminated orchards), the maximum of these surveillance frequencies is applied. Surveillance dates are drawn from a uniform distribution between the 92th and the 207th day of the year, i.e., the earliest and the latest surveillance dates in the database collected in southeastern France, respectively. If the cultivar produces flowers with petals (on which symptoms may be observed), the surveillance period is extended from the 59th to the 207th day.

#### Removals and replantings

Detected trees are removed individually and are not supposed to be replanted, because it would lead to orchard desynchronization in terms of phenology and fruit maturation. However, to avoid excessive fragmentation of orchards due to individual removals, orchards are totally removed if the proportion of living trees (i.e., trees in states S, E, H and D) falls below *χ_SEHD_*, which is a threshold for economic profitability (see details in supplementary Table S1). Moreover, detected trees trigger the definition of two nested zones (unrelated to the observation zones) around their orchard: all orchards in the removal zone (whose radius is *ζ_R_*) must be removed if the contamination rate *q_eR_* of a ‘removal epicentre’ (nested within the removal zone and defined by a radius *ζ_eR_*<*ζ_R_*), exceeds a threshold value *χ_R_*(Fig. 1D). Two Boolean parameters, *Y_R_^T^* and *Y_R_*, indicate whether individual detected trees or whole orchards, respectively, are to be removed after a mean delay of *δ*, or at the end of the year. Orchards can be replanted after a delay of *γ_S_* years. However, planting is forbidden if the contamination rate of the environment exceeds a threshold value 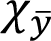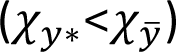, or if an orchard located at a distance below *ζ_n_* (this radius defines a zone called ‘neighbourhood’) has a contamination rate above a threshold *χ_n_* (Fig. 1C).

#### Output variables

The model computes two output variables: the net present value (*NPV*; in euros) of prunus cultivation, and the equivalent number of fully productive trees per hectare and per year (*Y*). The former is used as an economic optimisation criterion in our approach based on sensitivity analyses. Specifically designed for the present work, it is computed from the first (*a_m_*=6) to the last (*a_f_*=35) year of sharka management. Considering the whole production area as a single ‘farm’ with all the benefits and costs associated with prunus cultivation and sharka management, for year *a* (*a*=*a_m_*, …, *a_f_*) the gross margin (noted *GM_a_*) of this ‘farm’ can be calculated as the benefit generated by fruit sales (*p*; in euros/ha), minus fruit harvest costs (*c_H_*; in euros/ha), fixed cultivation costs (*c_F_*; in euros/ha), orchard planting cost (*c_S_*; in euros/ha), orchard observation cost (*c_O_*; in euros/ha), and the cost of removal of a tree (*c_R_^T^*; in euros) or a whole orchard (*c_R_*; in euros/ha). Reference values for these economic parameters were estimated (supplementary Table S1) based on expert opinions (i.e., professional organisations in charge of sharka management in France), as well as data on French peach production (Agreste 2013, 2014, 2015), selling price (FranceAgriMer 2015), and consumer price index (INSEE 2015). The cost of one orchard observation (*c_O_*) is described by a simple linear function of the detection probability: *c_O_* = 40 + 182 × *ρ*. This function is calibrated via expert opinions considering a fixed cost of access to the patch, and a surveillance cost proportional to the detection effort. This last feature allows to account for the effect of a partial surveillance of orchards (e.g., every other row only), which reduces both the average detection probability (*ρ*) and the observation cost. Thus, for all orchards *i* (*i* ∈ {1, …, *I*}) of the production area during year *a*, *GM_a_* is calculated as follows:

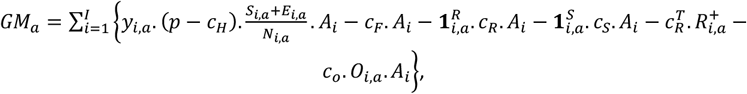

with:

*y_i,a_* the relative fruit yield of non-diseased trees depending on their age (see Table 1);
*A_i_* the orchard area (ha);
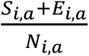 the proportion of the orchard that is not impacted by the disease;
*O_i,a_* the number of observations within the orchard;
*R^+^_i,a_* the number of newly (individually) removed trees due to PPV detection;
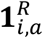 a Boolean which equals 1 if the orchard is removed, and 0 otherwise;
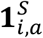 a Boolean which equals 1 if the orchard is planted, and 0 otherwise.

Using a discount rate *τ_a_*=4% (Quinet 2013), the net present value for the whole production area over years *a_m_*=6 to *a_f_*=35 is:

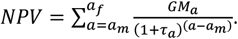

The equivalent number of fully productive trees (*Y*) was used to assess the epidemiological performance of management strategies that were economically improved using the *NPV*. This criterion was defined previously (Rimbaud et al. 2018a) by:

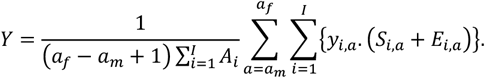

In this criterion, fully productive trees (i.e., mature and healthy) count for 1, whereas diseased and newly planted trees (juvenile) count for 0.

### Management improvement through sequential sensitivity analyses

Improvement of the management strategy was performed using three sensitivity analyses. For these analyses, we used Sobol’s method, which is a reference method to compute sensitivity indices of model parameters and their interactions (Saltelli et al. 2008; Sobol 1993). Each sensitivity analysis consists of: i) defining the target parameters, their respective variation ranges and probability distributions (uniform, in this work); ii) generating a design to explore the parameter space (using Sobol’s sequences in this work; see details in Rimbaud et al. 2018a); iii) running simulations; and iv) computing Sobol’s sensitivity indices which quantify the influence of the variation of each target parameter on the output variable. Let *Y* denote the output variable, and *X_i_* (*i*=1,…*p*) the input parameters of the model. Sobol’s indices are calculated for each *X_i_* as follows:

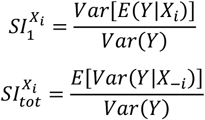

where *X_-i_* denotes the whole set of parameters except 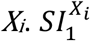 is the 1^st^-order index, which measures the main effect of *X_i_* alone; and 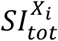 is the total index, which measures the influence of *X_i_* including all its interactions with other parameters. These dimensionless indices are bounded by 0 and 1, and a total index close to 0 means that the variation of the parameter has a negligible effect on the output variable.

To enable comparison among the sensitivity indices of the model parameters, Sobol’s method requires to sample the target parameters from independent distributions. Thus, since the focal zone (radius *ζ_f_*), the observation epicentre (*ζ_eO_*) and the removal epicentre (*ζ_eR_*) are contained within the security zone (*ζ_S_*), the focal zone (*ζ_f_*) and the removal zone (*ζ_R_*), respectively, these zones were re-parameterised using the following area ratios (bounded by 0 and 1):

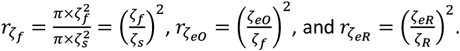

In addition, the contamination threshold above which surveillance frequency in young orchards is modified (*χ_y*_*), was re-parameterised using its ratio relative to the contamination threshold above which replanting is forbidden 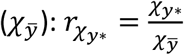.

#### Step 1: Assessing the relative influence of model parameters

The 23 control parameters defined in the implemented management strategy were targeted in this first step, in addition to 6 parameters associated with the main epidemiological processes (introduction: *q_κ_*, *Φ*, and *pMI*; dispersal: *W_exp_*; transmission coefficient: *β*; and latent period duration: *θ_exp_*). Except for *q_κ_* (connectivity of the patch of first introduction) and *p_MI_* (probability of massive introduction), variation ranges of the epidemiological parameters were defined as the 99% credibility intervals of the estimates provided by a Bayesian inference model applied to PPV-M epidemics in southeastern France (Pleydell et al. 2018). In contrast, variation ranges of *q_κ_*, *PMI* and the 23 control parameters were informed by expert opinions and restricted to realistic values with respect to sharka epidemics and management (Table 1).

Simulations were performed for 310,155 different parameter combinations generated with Sobol sequences (Sobol 1976). To account for stochasticity (see, e.g., Rimbaud et al. 2018a), each combination was replicated 30 times (this number, above which the total computational time would be prohibitive, was sufficient to obtain robust estimates of the mean and standard deviation associated with each combination; supplementary Fig. S1). Then, the indices were calculated as in (Rimbaud et al. 2018a) to assess the influence of target parameters on the means and standard deviations of the economic (*μNPV* and *σNPV*) and epidemiological (*μY* and *σY*) criteria.

#### Step 2: Approximating optimal values of the most influential parameters

In the second step, only the most influential control parameters (i.e., those with the highest *SI_tot_*) identified in the first step were retained (see Results). The 6 least influential control parameters were removed from the model, and the 17 remaining parameters were varied within the same variation ranges as previously (Table 1), using 310,156 different parameter combinations and 30 stochastic replicates. In this sensitivity analysis, the 6 epidemiological parameters were not targeted, but for each simulation they were drawn from uniform distributions using the same bounds as in the previous step, in order to optimise the management strategy for variable (but realistic) epidemics.

The optimal values of the 17 control parameters were jointly approximated by identifying the parameter combination associated with the highest *μNPV* (‘best-value strategy’). To identify alternative improved strategies, a marginal approximation of the same parameters was also performed, using the mode of the distribution of each parameter within the combinations associated with the best 1% values of *μNPV* (‘best-percent strategy’). A similar approach allowed identification of the parameters associated with the worst 1% values of *μNPV*. The marginal approximation (performed parameter by parameter) does not account for possible interactions between parameters, contrary to the first method that retains a whole parameter combination.

#### Step 3: Improving the approximate optimal values

Based on the results of the previous step, six control parameters were found unnecessary (see Results) and subsequently removed from the model. One control parameter had the same value for all strategies with the best *NPVs*, and was thus fixed at this value. The optimal values of the 10 remaining control parameters were still very imprecise; thus, these 10 parameters were further improved using a dedicated sensitivity analysis with variation ranges restricted to the intervals where most of the best 1% values of *μNPV* were found in step 2 (Table 1). This sensitivity analysis was performed as in previous steps, using 310,152 different parameter combinations and 30 replicates. For each simulated epidemic, the epidemiological parameters were drawn from uniform distributions as in step 2.

### Simulation of the improved management strategies

To test the performance of the identified management strategies, 10,000 simulation replicates were performed with parameters corresponding to different scenarios: (A) “Disease-free” (PPV is not introduced in the landscape); (B) “Management-free” (PPV is introduced and not managed); (C) “Reference management” (PPV is introduced and managed according to the current French strategy; Table 1); (D) “best-value strategy” and (E) “best-percent strategy” (Table 2). In scenarios B to E, the six epidemiological parameters were drawn from uniform distributions within their respective variation ranges (Table 1). The impact of the epidemic was assessed using the distribution of the epidemiological criterion (*Y*), the economic criterion (*NPV*), as well as the dynamics of annual prevalence and incidence, and the dynamics of observations and removals.

**TABLE 2.**
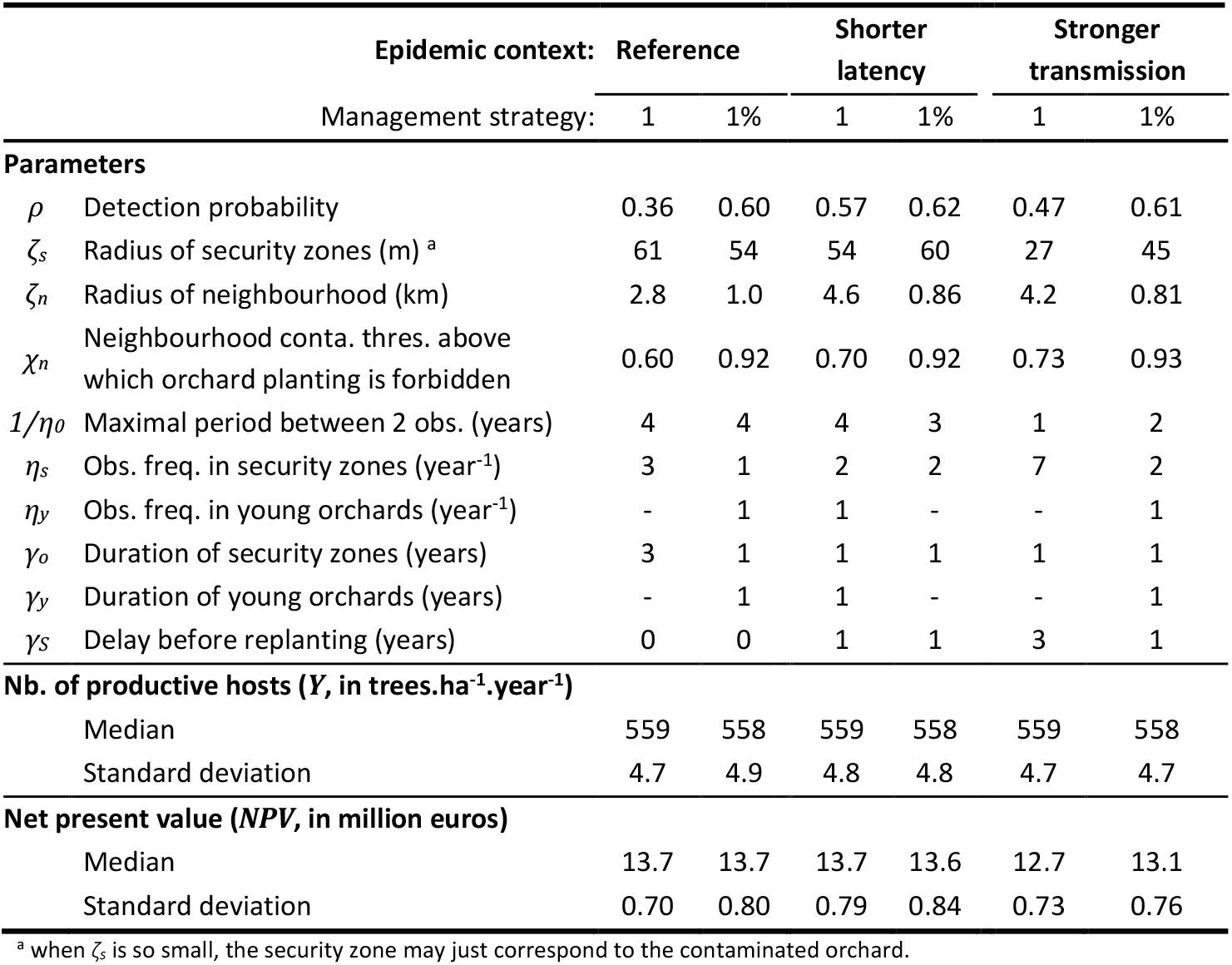
Summary of the economically improved management strategies in different epidemic contexts. Values of the control parameters estimated using the combination leading to the best value (1), or the highest percentile (1%) of *μNPV*, and outputs of 10,000 simulations, in the reference epidemic context or in harsh epidemic contexts (halved values for the expected duration of the latent period, *θ_exp_*; or doubled values for the transmission coefficient, *β*).

Furthermore, the robustness of the identified management strategies to harsher epidemic contexts was tested using similar simulations with either doubled values for the pathogen transmission coefficient (*β*) or halved values for the latent period duration (*θ_exp_*). New strategies, specifically improved for each context, were identified by following the three steps of our heuristic approach.

### Computing tools

The model was written in R and C languages; one simulation takes around 8 s on an Intel^®^ Core™ i7-4600M computer. Within the R software v3.0.3 (R Core Team 2012), Sobol’s sequences were generated and Sobol’s indices were calculated using the packages *fOptions* v3010.83 (Wuertz et al. 2017) and *sensitivity* v1.11 (Pujol et al. 2017), respectively.

## RESULTS

We identified improved management strategies of sharka epidemics in three steps, each of them consisting of a sensitivity analysis of a model which jointly simulates sharka epidemics and a flexible management strategy (see supplementary Fig. S2 for a general illustration).

### Step 1: Identifying the most promising control parameters

In a first step, the most promising control measures to manage sharka were identified by ranking the control parameters by influence on the economic criterion (average net present value over 30 years, *μNPV*, measuring the discounted difference between benefits of fruit sales and costs generated by the management of all orchards of the production area). The first sensitivity analysis showed that the predicted economic impact strongly depends (positively; see Fig. 2) on the contamination threshold of the ‘removal epicentre’ (*χ_R_* above which orchards in the removal zone are culled; total sensitivity index, *SI_tot_*: 0.54; CI_95_: 0.39-0.65), and on the contamination threshold in orchards of the neighbourhood of a previously culled orchard (*χ_n_*, above which replanting is forbidden; *SI_tot_*: 0.36; CI_95_: 0.25-0.45). These parameters were also the main contributors to model stochasticity (supplementary Fig. S3). Next comes the radius of the removal zone (*ζ_R_*; *SI_tot_*: 0.15; CI_95_: 0.08-0.20), then a Boolean parameter indicating whether whole orchards are removed immediately or at the end of the year (*ϒ_R_*; *SI_tot_*: 0.13, CI_95_: 0.06-0.18), followed by the detection probability (*ρ*; *SI_tot_*: 0.09; CI_95_: 0.07-0.12) and the size of the removal epicentre–where the contamination rate is measured to assess if orchards in the removal zone (with a radius *ζ_R_*) must be culled (*r_ζ_eR__*; *SI_tot_*: 0.09; CI_95_: 0.05-0.12) (Fig. 2). The effect of each of these parameters seems to be mostly due to interactions with other parameters, since their total Sobol indices were much higher than their 1^st^-order indices. The importance of these interactions explains why it is difficult to predict the net present value (*NPV*) from the values of *ζ_R_*, *ϒ_R_* or *r_ζ_eR__* alone (Fig. 2, insets). The strong impact of removal and planting measures on the mean *NPV*(*μNPV*) is likely due to excessive tree removals or orchard planting bans when the respective contamination thresholds are low (below 5% for removals and below 10% for planting bans; Fig. 2, insets). However, for higher thresholds, *μNPV* remained stable, suggesting that these parameters no longer have an effect. This may be the sign of the absence (or rarity) of high contamination events in orchards or in removal epicentres (and thus of extremely rare planting bans or removal of whole zones). Finally, the quadratic effect of *ρ* reveals the existence of a trade-off between detection efficiency and the costs induced by orchard surveillance.

**Fig. 2.**
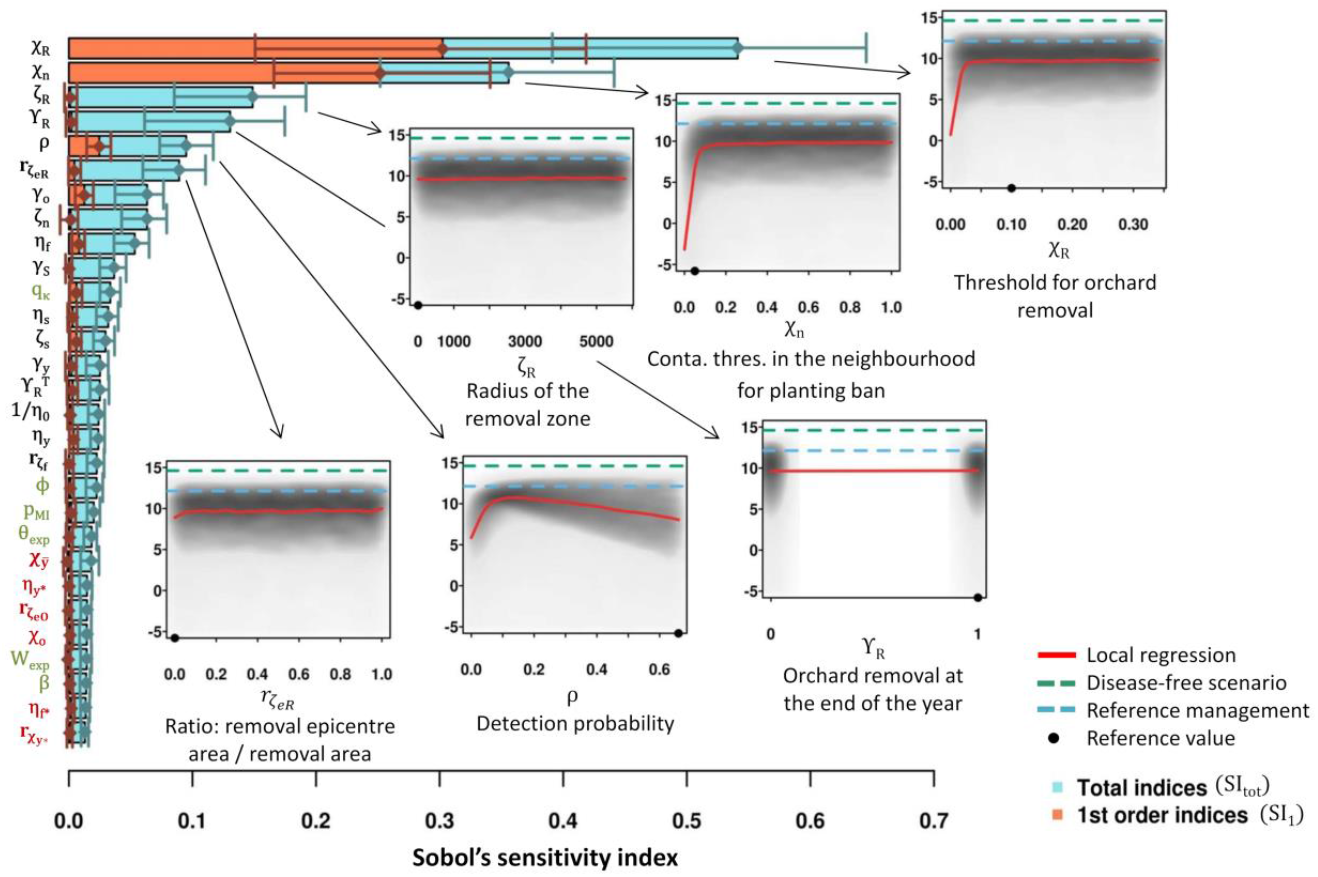
Step 1: Sobol’s sensitivity indices of the 23 control parameters and 6 epidemiological parameters on the mean output of 30 stochastic replicates (*μNPV*, average net present value). 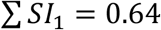. Parameters in black are kept in step 2; parameters in red are removed; parameters in green are the epidemiological parameters. Insets: value of the output variable (*μNPV*, in million €) obtained with the different values of the six most influential parameters (grey colouration: density). All model parameters are defined in Table 1.

The 11 least influential parameters (5 epidemiological parameters and 6 control parameters) had very low total sensitivity indices (less than 0.03), which indicates a negligible effect of their variation on the mean output. These 11 parameters included all epidemiological parameters except the connectivity of the patch of first pathogen introduction (*q_κ_*). This suggests that, in the simulated context, the economic impact of epidemics depends more on control actions than on the epidemics themselves (except for the location of the first PPV introduction). Nevertheless, some control actions (those governed by the 6 least influential control parameters) also have little effect on the economic impact of epidemics. These control actions relate to the reinforcement of visual surveillance in locally highly contaminated areas (*r_ζ_eO__*, *χ_O_* and *η_f*_*) or in young orchards (*r_χ_y*__* and *η_y*_*), or to planting bans after orchard removals (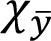). The negligible influence of these control actions can be attributed to the fact that in the simulations, the level of local contamination never reached the thresholds for surveillance reinforcement or planting ban. Similar conclusions emerged from the analysis of the epidemiological output of the model (number of fully productive trees per hectare and per year, *μ_Y_*; supplementary Fig. S4). Consequently, the 6 least influential control parameters were removed in the next step in order to more efficiently improve sharka management strategy.

### Step 2: Approximating optimal values of control parameters

A second sensitivity analysis was performed on the control parameters only (except for the 6 control parameters discarded in step 1). Optimal values of the 17 control parameters were approximated either by the parameter combination leading to the best *μNPV* (‘best-value strategy’), or the respective modes of the parameter distributions leading to the highest percentile of *μNPV* (‘best-percent strategy’). In both approaches, management is improved when detected trees are removed immediately (i.e., after an average of 10 days, rather than at the end of the year; see *Y_R_^T^* in Fig. 3). The obtained values for the removal threshold (*χ_R_*, mostly above 5%), the radius of the removal zone (*ζ_R_*, mostly above 1000 m), and its epicentre where the contamination rate is measured (*r_ζ_eR__*, mostly above 0.1), were so high that no removal of whole orchards may occur in the associated simulations (see *χ_R_* in Fig. 3 and all other parameters in supplementary Fig. S5). In addition, small values for these parameters (*χ_R_* < 5%; *ζ_R_* < 2 km; *r_ζ_eR__* < 0.2) were associated with the lowest percentile of *μNPV*. These results suggest that in our model, the removal of healthy hosts induces higher costs than the maintenance of undetected or latently infected hosts. Regarding surveillance, either the focal area matched approximately the security area (*r_ζ_f__* and *ζ_s_* in the ‘best-value strategy’), or surveillance frequencies in these zones were the same (*η_f_* and *η_s_* in the ‘best-percent strategy’). This indicates that it may not be necessary to distinguish between focal and security zones. Thus, parameters associated with removal zones (*χ_R_*, *ζ_R_*, *r_ζ_eR__* and *ϒ_R_*), and focal zones (*χ_ζ_f__* and *η_f_*) were discarded in the next step.

**Fig. 3.**
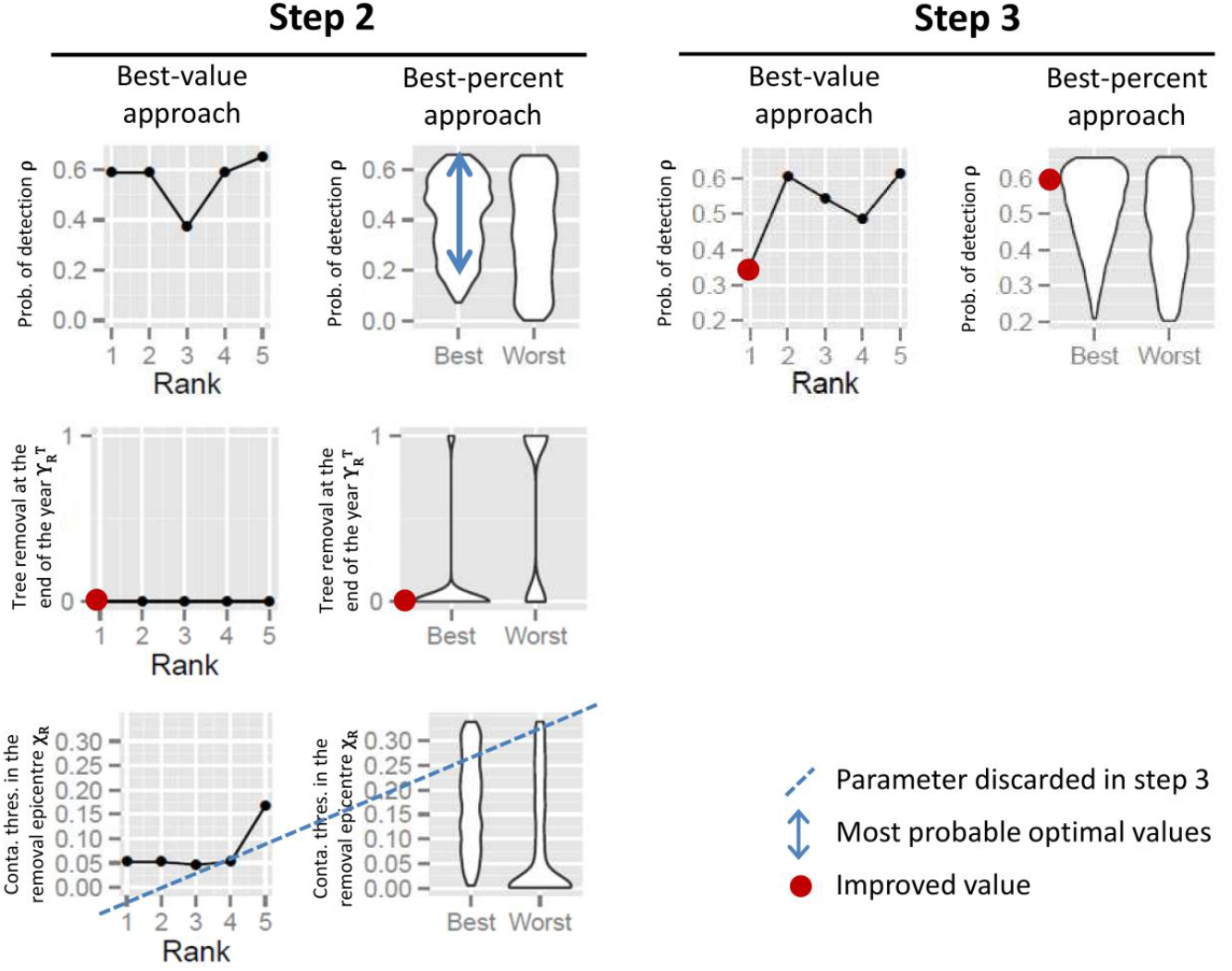
Approximation of the optimal control parameters through sequential sensitivity analyses: example on three parameters. We use the combination associated with the highest value of *μNPV*, or the mode of the distribution of each parameter within the combinations associated with the highest percentile of *μNPV*. In step 2, an improved value (red circle) is found for *ϒ_R^T^_*, and *χ_R_* can be removed from further analyses (dashed blue line: improved values for removal parameters are so high that removals may never occur). The approximation of *ρ* is improved in step 3, using a restricted variation range (blue arrows). See the results of steps 2 and 3 with all control parameters in supplementary Figs. S5 and S6, respectively.

### Step 3: Improving the approximation

A third sensitivity analysis was performed on the 10 remaining control parameters to further refine the approximation of their optimal value. In this analysis, variation ranges of the target parameters were restricted to the intervals where most of the best 1% values of *μNPV* were found in the previous step (see the probability of detection *ρ* in Fig. 3 and all other parameters in supplementary Fig. S5). Then, optimal values were approximated as in step 2 (Fig. 3 and supplementary Fig. S6). The resulting ‘best-value’ and ‘best-percent’ strategies are significantly more cost-effective and simpler than the current French strategy (Fig. 4). To summarise the best-value strategy, every time a symptomatic tree is detected in an orchard, this orchard is visually surveyed three times a year for the three following years (Table 2, reference epidemic context). Whole orchards are removed only if the proportion of living trees falls below *χ_SEHD_*=0.66 (for profitability reasons, see Materials and Methods, and supplementary Table S1), and in this case orchards can be replanted without delay. Nevertheless, replanting is forbidden if the contamination rate of an orchard exceeds 60% within a radius of 2.8 km (i.e., in case of a massive introduction, in our simulations). In areas where no infected tree has been detected in the three previous years, orchards are surveyed once every four years. The detection probability is 36%, i.e., nearly half the reference value (66%), which in practice means that only every other row is surveyed in the orchards.

**Fig. 4.**
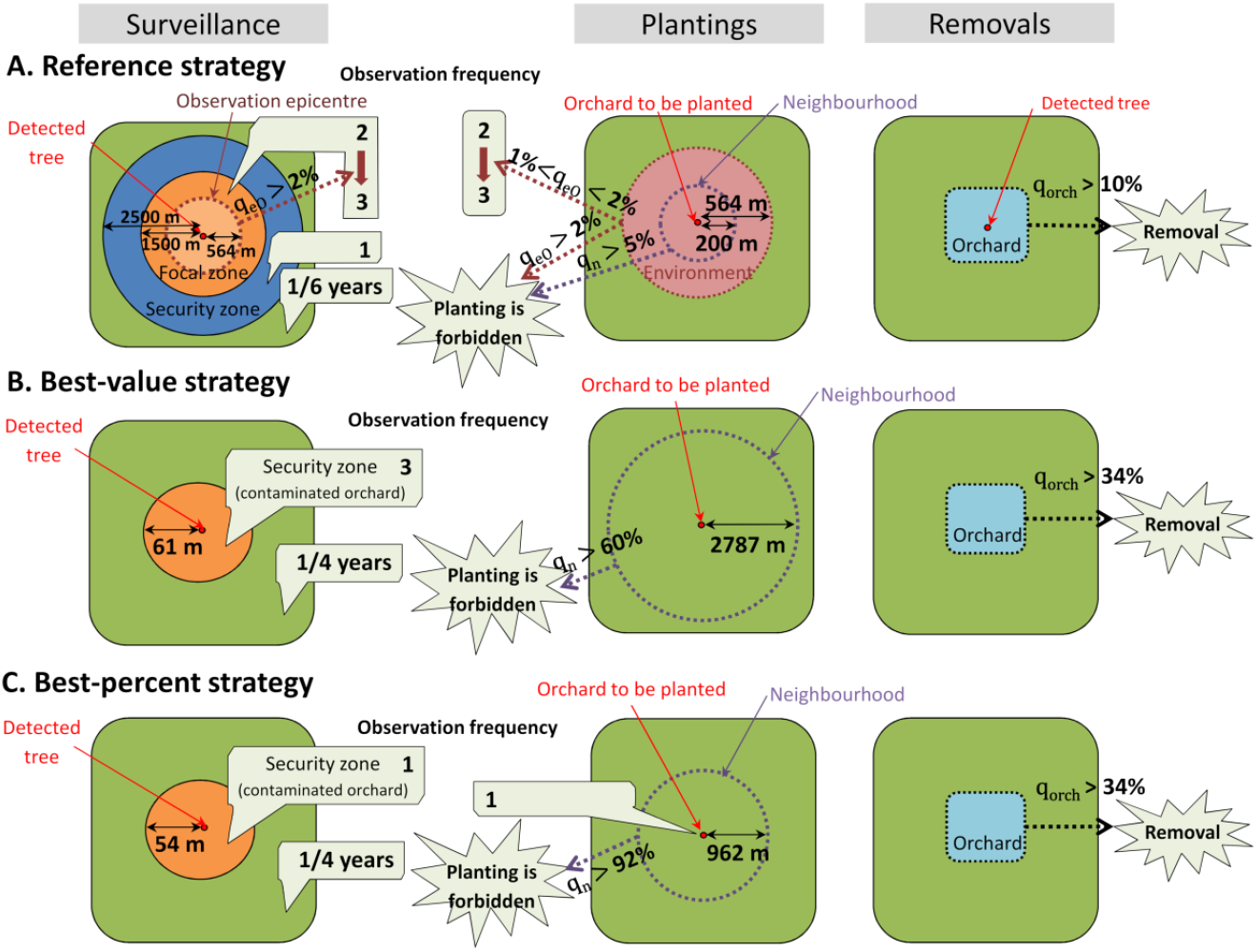
Management strategies. Surveillance, plantings and removals according to the reference management (French management in orchards; A), or to the combination of control parameters associated with best value (B) or the highest percentile (C) of the economic criterion *μNPV*. In panels B and C, given its radius, the security zone may consist of the contaminated orchard only; whole orchards may only be removed if the proportion of remaining trees is below the profitability threshold (*χ_SEHD_*=0.66).

Alternatively, in the best-percent strategy (Table 2), each time a symptomatic tree is detected, the corresponding orchard is surveyed once in the following year. Similar to the best-value strategy, whole orchards may only be removed for profitability reasons, and can be replanted without delay (except in the unlikely event of a contamination rate higher than 92% in an orchard within a radius of 962 m). In the first year after planting, young orchards are surveyed once. Older orchards in non-contaminated areas are routinely surveyed every four years. The detection probability is 60% (i.e., close to the reference value).

### Simulation of the improved strategies

Repeated stochastic simulations were carried out with the identified improved strategies in order to compare their epidemiological (*Y*) and economic (*NPV*) outcomes with those obtained in disease-free and management-free scenarios, and with the reference management (i.e., the current strategy for managing sharka in France). First, these simulations show that managing the disease is vital, since the median number of productive trees per hectare and per year in the management-free scenario is only 513 and is highly variable [2.5%; 97.5% quantiles: 462; 551] compared to the disease-free scenario (median number of 560 trees [551; 569]) (Fig. 5a). This results in a 59% decrease in the median *NPV* (6.03 million euros [-4.06; 13.0] in the absence of management, *vs* 14.6 million euros [13.5; 15.7] in the absence of disease), and possibly huge economic losses in the worst-case scenarios (Fig. 5B). In our model, the reference strategy is very efficient to reduce the epidemiological impact of sharka (median of 559 productive trees per ha and per year [550; 569]). The associated *NPV* (12.1 million euros [10.7; 13.4]) is much higher than in the absence of management; however, it is 17% lower than in the disease-free scenario, indicating considerable costs of disease management. The improved best-value and best-percent strategies are nearly as efficient as the reference strategy with regard to the number of productive trees, but resulted in reduced costs, with a median *NPV* of 13.7 [12.2; 15.0] and 13.7 [11.7; 15.0] million euros, respectively. These management strategies are associated with slightly higher disease incidence and prevalence than for the current French strategy (supplementary Fig. S7, top row). The resulting increase in individual tree removal is overly compensated by the smaller number of observations in orchards (supplementary Fig. S8, top row). Consequently, both improved strategies are more cost-effective than the reference strategy.

**Fig. 5.**
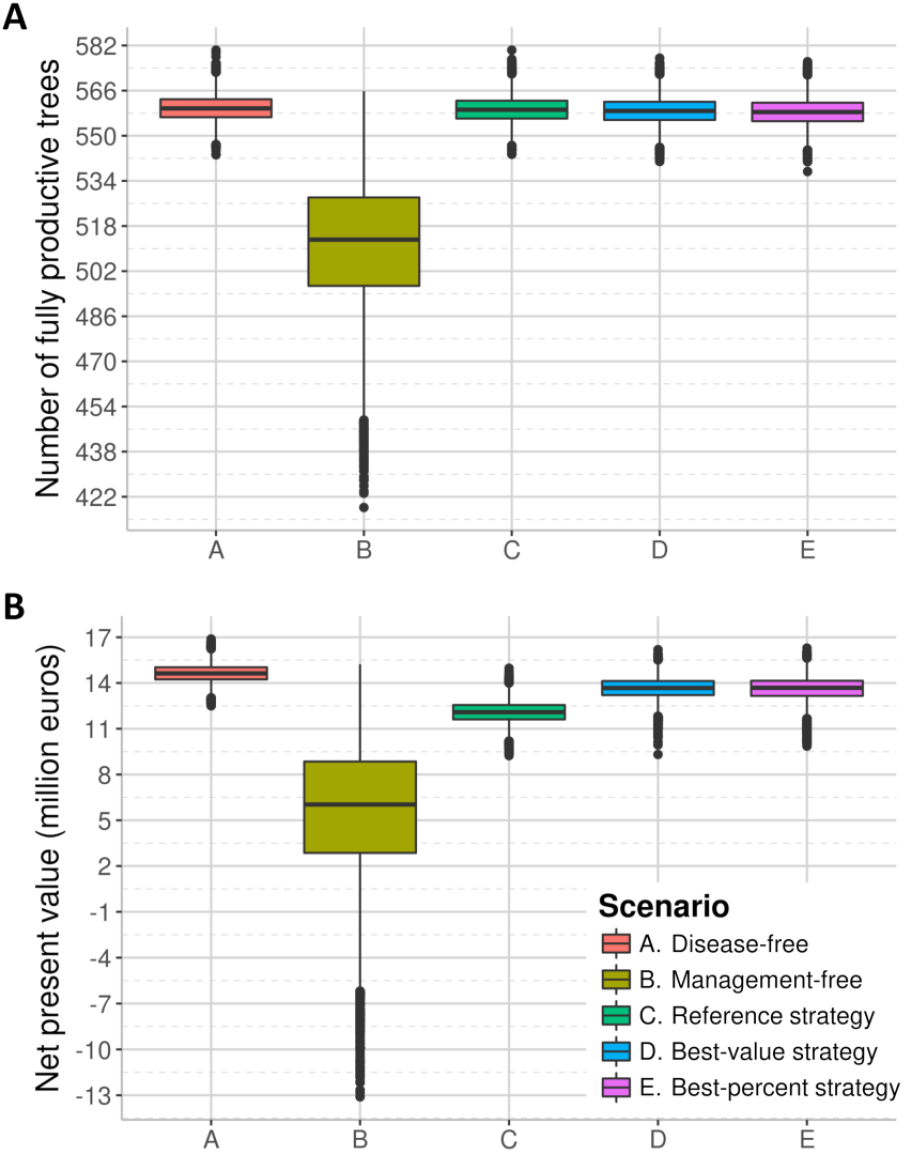
Boxplots of *Y* (equivalent number of fully productive trees per hectare and per year, A), and *NPV* (net present value of all orchards of the landscape, B) after 30 years of sharka management. Different scenarios are simulated: absence of disease, absence of management, disease managed with the reference strategy (French management in orchards), or with economically improved management strategies identified through two different methods (combination associated with the best value or the highest percentile of *μNPV*).

When tested in harsher epidemiological contexts with halved expected duration of the latent period of the pathogen (*θ*_exp_), the identified management strategies were still efficient with respect to both the epidemiological (*Y*) and economic (*NPV*) criteria (supplementary Fig. S9, panels A and B). However, with doubled transmission coefficient (*β*), the improved management strategies (in particular the best-percent strategy) were not as efficient as in the reference scenario (supplementary Fig. S9, panels C and D). This resulted in higher levels of disease prevalence and incidence (supplementary Fig. S7, bottom row) and consequently more tree removals (supplementary Fig. S8, bottom row). In contrast, the reference management strategy was robust to the simulated changes in epidemic severity (supplementary Figs. S7, S8 and S9).

To specifically improve the management strategy in harsher epidemic contexts, we followed again all the steps of the proposed approach. With a shorter latent period of the pathogen, the newly identified strategies mainly differ from the previous ones by the introduction of a 1-year delay before orchard replanting. This delay, which matches the mean duration of the latent period (*θ_exp_*=0.96 year in this scenario), enables the removal of infected trees in the neighbourhood before planting a new orchard (Table 2). With stronger transmission coefficient (*β*=1.64), the most striking difference with the reference context is in the ‘best-value’ strategy, which relies on a 3-year delay before replanting, and strengthened surveillance frequencies: all orchards are surveyed every year, and those located in security zones are surveyed 7 times a year (Table 2). These measures allow disease suppression despite its faster expansion. As shown with the repeated simulations in harsh epidemic contexts, these new strategies satisfy both epidemiological and economic performance criteria (supplementary Figs. S10 and S11).

## DISCUSSION

### A method to improve management strategies of epidemics

In this article, we propose an innovative heuristic approach based on the sequential use of global sensitivity analysis of a simulation model to improve strategies for the management of plant disease epidemics. Although parameter ranges might have been restricted to likely values using formal (Bayesian) elicitation of expert’s opinion, we chose to start from the widest meaningful ranges and narrow them down at each step of the procedure. Following the identification of the most promising control measures (step 1), the exploration of parameter space, initially large (step 2) and then more restricted (step 3), enables to simplify the strategy and to progressively improve the control parameters (supplementary Fig. S2). This method is based on the identification of the best strategies among a finite set of simulations. Consequently, the identified strategies are only approximately optimal, despite the wide range of explored scenarios (further sensitivity analyses could be performed if more accuracy was required for some parameters). Nevertheless, in contrast with previous studies focussing on a restricted number of control parameters (e.g., Cunniffe et al. 2015; Hyatt-Twynam et al. 2017; Parnell et al. 2010), our approach includes a wide range of management strategies and enables to jointly improve the entire set of control parameters. Point estimates of improved control parameters were identified using two different approaches (‘best-value’ and ‘best-percent’) resulting in two specific options to improve the reference management strategy (Fig. 4, panels B and C). Alternatively, it could be possible to extract, from the sensitivity analyses, whole distributions of parameter values leading to the best percentile of *μNPV* (supplementary Fig. S6). While this method would not account for potential interactions between parameters, it would inform on the degree of precision required on each control parameter to apply the management strategy.

### Balancing costs and benefits of disease management

The selection of a relevant criterion is a crucial task in an optimisation procedure. Epidemiological criteria that have been minimised for management optimisation include disease incidence (Breukers et al. 2007; Holt et al. 1999; Xu and Ridout 1998), prevalence (Courcoul et al. 2011; Lurette et al. 2009), severity (Xu and Ridout 1998), propagation rate (Coutts et al. 2011; Filipe et al. 2012), basic reproduction number (Chan and Jeger 1994), proportion of infected plants at the end of the simulation (Sisterson and Stenger 2013), probability of pathogen establishment (Papaïx et al. 2014), time until eradication (Barclay and Vreysen 2011; Courcoul et al. 2011; Jeger and Chan 1995; Parnell et al. 2009, 2010), or proportion/number of removed trees required to achieve eradication (Hyatt-Twynam et al. 2017; Parnell et al. 2009). However, to be feasible and sustainable, the choice of a management strategy must be guided by the balance between the costs induced by control actions and the economic benefits of the reduction of epidemic damage (Forster and Gilligan 2007; Fraser et al. 2004). Some studies proposed approaches to prioritise surveillance or removal options under limited resources (Cunniffe et al. 2016; Faulkner et al. 2016; Ndeffo Mbah and Gilligan 2010). Other studies proposed optimised management of different plant, animal and human diseases accounting for the cost of management (e.g., surveillance and removal, culling, vaccination) relative to the intrinsic cost of the epidemic or the benefit of maintaining a healthy population (Cunniffe et al. 2014; Forster and Gilligan 2007; Kleczkowski et al. 2012; Thompson et al. 2018). Here, we used an economic criterion, the net present value (*NPV*), which explicitly accounts for the benefits (i.e., fruit sales) generated by the cultivation of productive hosts, and the costs induced by orchard cultivation plus different control actions comprising surveillance, removal, replanting, and planting bans. By doing so, the different components of the management strategy can be optimised with explicit consideration of their cost relative to the benefit due to the reduction of epidemic damage. In addition, the inclusion of a discount rate—a key element in project financial evaluation—accounts for the time value of money, i.e., the fact that the same income has less value in the future than in the present. For these reasons, the *NPV* is a powerful criterion to identify cost-effective strategies for the management of epidemics. Although this criterion was specifically parameterised for peach orchards in this work, it could easily be employed for other crops by reassessing the costs and benefits associated with their management. Finally, we also used an epidemiological criterion to assess the epidemiological efficiency of the identified improved strategies. This metric integrated the number of productive hosts across the whole production area and the whole simulation period, and is therefore comparable to the ‘healthy area duration’ used in many epidemiological models (e.g., Lo Iacono et al. 2013; Papaïx et al. 2018; Rimbaud et al. 2018b). By using this metric, we aimed to maximise the number of productive hosts rather than to minimise the number of infected hosts, because the latter option does not penalise the removal of whole orchards and their replacement by young (non-productive) trees.

### Improving sharka management

#### Identifying promising control measures

Based on Sobol’s indices, the first sensitivity analysis highlighted the most influential control parameters on sharka epidemics in French peach orchards. These were associated with orchard removals, planting bans, and detection probability (Fig. 2). Although Sobol’s method only indicates the relative influence of the parameters and not the direction of change in the output variable, it can be complemented by graphical methods or metamodels (Rimbaud et al. 2018a). Here, the smoothed curves showed the existence of an optimal detection probability, and a positive correlation between the *NPV* and the threshold values for removal and planting bans (Fig. 2, insets). These elements highlight the importance of modelling studies designed to optimise detection efficiency (e.g., using hierarchical sampling; Hughes et al. 1997) and removal actions (e.g., by identifying an optimal cull radius; Cunniffe et al. 2015, 2016; Parnell et al. 2009, 2010).

Ranking model parameters by their influence on a given output variable also allows the identification of weak contributors. Among the implemented control parameters, those associated with the modification of surveillance in response to the observed local prevalence had a negligible impact on both epidemiological and economic outcomes (Fig. 2). Therefore, in the investigated scenarios, reinforcing surveillance in highly contaminated areas may not result in improved management. Thus, the associated parameters may be discarded from the modelled management strategy without altering its efficiency. However, we assumed that each symptomatic tree had the same detection probability, regardless of its age, cultivar and time elapsed since symptom expression. Nevertheless, field experience shows that all these factors impact symptom expression and consequently visual detection. Moreover, detection events were considered independent, which enabled an excellent global detection rate after a few surveillance rounds. All these elements could partly explain why reinforcing surveillance in highly contaminated areas had a negligible impact in our analysis, but might be useful in the field.

In previous studies, the connectivity of the patch of first introduction was the most influential epidemiological parameter on sharka spread in the absence of management (Rimbaud et al. 2018a). It is still the case here, in the presence of management (Fig. 2, parameter *q_κ_*). This result suggests that reinforced baseline surveillance of highly connected patches could be a promising control measure. This result also supports the need to match management efforts with the risk index of different patches (Barnes et al. 1999; Nelson et al. 1994; Parnell et al. 2014) or individual hosts (Hyatt-Twynam et al. 2017) and to prioritise such management under limited resources (Cunniffe et al. 2016; Faulkner et al. 2016). Favouring the planting of resistant cultivars (provided that they are available) in patches with high risk indices could also be a promising approach for disease management.

#### Identifying improved strategies

Our results suggest that removing whole orchards triggers higher costs than potential losses associated with maintaining undetected hosts (i.e., infectious trees in state H). The identified improved strategies (targeting a cost-effective management rather than eradication) are thus based on a ‘symptomatic hosts hunt’ (Fig. 4, panels B and C). However, orchard surveillance to identify these symptomatic hosts also induces high costs. Thus, in contaminated orchards, improved strategies relied either on few surveillance rounds with a good probability of detection (close to the reference value), or a greater number of surveillance rounds with a low probability of detection (half the reference value, which may correspond to the surveillance of only every other row in the orchard to reduce the cost of a surveillance round). Overall, while surveying an orchard more than once a year multiplies the costs, it favours the detection of infected trees which may remain undetected after the first round of surveillance. It might be interesting to investigate the potential of alternative detection protocols, like serological or molecular methods, which are used for the management of several perennial plant epidemics (including sharka in the United States; Gougherty et al. 2015). Such investigation may be performed with our model by changing the relationship between the probability of detection and the cost of surveillance.

Our results indicate that the current French management of sharka should already be very efficient in controlling the disease (Fig. 5A); consequently, no major improvement should be expected with respect to the epidemiological criterion. In contrast, the economic criterion shows that the French strategy is associated with substantial costs (Fig. 5B). We identified improved strategies, which should be epidemiologically as efficient as the current French strategy but less expensive–and also simpler and hence easier to implement in the field. These strategies are based on simplified surveillance procedures, and on the absence of removal zones (Fig. 4). Nevertheless, the default surveillance frequency (for orchards located outside of focal and security zones) is increased to once every 4 (instead of 6) years. This change enables to remove newly introduced infectious trees more promptly after the end of the latent period, which was shown to decrease the overall management costs (Cunniffe et al. 2016).

Several assumptions may influence the improved strategies identified here. In particular, the ability to detect symptomatic hosts may have been overestimated as explained above, which should result in practice in a slight under-control. Consequently, in the field and under certain conditions, it might still be interesting to rogue whole orchards in order to remove hard-to-detect diseased trees. Furthermore, we assumed that all orchards had the same susceptibility to PPV. However, young plants may be more susceptible than older ones (Astier et al. 2007). Thus, our results may underestimate the importance of surveying young orchards.

Despite these considerations, the best-value and best-percent strategies deserve to be considered in order to improve the cost-efficiency of sharka management. The common feature of these options is a simplification of surveillance and removal measures that reduces management costs while maintaining good epidemiological performance. However, our results show that these strategies may fail to control disease in harsh epidemic contexts. For these situations, the flexibility of our approach allowed us to identify new strategies, illustrating its general ability to find improved combinations of control parameters in a given epidemic context. Furthermore, such strategies are relevant to the other (perennial) plant diseases that are managed through surveillance, removals and planting restrictions. We note, however, that some control actions (for which we had no cost/benefit values) were not investigated in this work, like nursery protection measures, planting at different densities, or the use of resistant prunus trees. In addition, vector control by insecticides was not studied here because insecticides generally fail to control the spread of non-persistent viruses such as PPV (Perring et al. 1999; Rimbaud et al. 2015).

## ACKNOWLEDGEMENTS

The authors thank G. Labonne and S. Grizard for constructing the surveillance database, SRAL RA, FREDON 30, and especially J. Aymard for their expertise in prunus cultivation, D.R.J. Pleydell for programming the first version of the model, P.H. Thrall for reviewing a previous version of this manuscript, the reviewers and Editor for their valuable comments, the INRA BioSP and the CIRAD SouthGreen bioinformatics platforms and L. Houde and S. Ravel for providing support.

## SUPPORTING INFORMATION LEGENDS

**Fig. S1**. Distribution of 5,000 means (A) and standard deviations (B) of the model output (net present value, *NPV*) computed with an increasing number of replicates. All model parameters are fixed at their reference value. The dashed vertical line delimits the selected number of replicates (30).

**Fig. S2**. Schematic representation of the heuristic approach to improve sharka management using sequential sensitivity analyses of a simulation model. In every sensitivity analysis (SA), variation ranges of epidemiological parameters are restricted to simulate realistic epidemics, whereas control parameters are varied within values that are compatible with a feasible management. In step 1, a first SA highlights key epidemiological and control parameters. In step 2, a second SA targets the most influential control parameters and approximates optimal values. Then, parameters associated with a wide range of improved values are further improved via a dedicated SA in step 3. Control actions governed by parameters with negligible influence (in step 1) or associated with unrealistic values (in step 2) can be removed from the model. Blue ellipses: sensitivity analyses; grey numbers: number of parameters in the model; boxes: outcomes of the approach.

**Fig. S3**. Step 1: Sobol’s sensitivity indices of the 23 control parameters and 6 epidemiological parameters on the standard deviation of the stochastic replicates (A: *σ_Y_*, average number of fully productive trees; B: *σ_NPV_*, average net present value). Parameters in black are kept in step 2; parameters in red are removed; parameters in green are the epidemiological parameters.

**Fig. S4**. Step 1: Sobol’s sensitivity indices of the 23 control parameters and 6 epidemiological parameters on the mean epidemiological output of the stochastic replicates (*μ_Y_*, average number of fully productive trees). Parameters in black are kept in step 2; parameters in red are removed; parameters in green are the epidemiological parameters. Results on the economic output (average net present value, *μNPV*) are presented in Fig. 2.

**Fig. S5**. Step 2: Five best values (first and third columns), and distribution of each parameter for the highest and lowest percentiles (second and last columns) of the economic criterion (*μNPV*, average net present value) in the second sensitivity analysis. A single improved value is found for 1 parameter (red circle), 6 unnecessary parameters can be removed (dashed red lines: the focal zone matches the security zone, so both zones can be merged; dashed blue lines: improved values for removal parameters are so high that removals might never occur) and the 10 remaining parameters are to be further improved in step 3, using restricted variation ranges (blue arrows).

**Fig. S6**. Step 3: Five best values (first and third columns), and distribution of each parameter for the highest and lowest percentiles (second and last columns) of the economic criterion (*μNPV*, average net present value) in the third sensitivity analysis. Improved values for each control parameter are indicated by red dots. The duration of young orchards (*γ_y_*) is irrelevant in the best-value strategy because the observation frequency in young orchards (*η_y_*) is 0.

**Fig. S7**. Dynamics of annual prevalence (blue curves) and incidence (red curves) under different management strategies (French management in orchards, or economically improved strategies) and epidemic contexts (reference context, or harsher epidemics: half the duration of the expected latent period, *θ_exp_*, or doubled transmission coefficient, *β*) in 100 simulations. The dashed vertical line indicates the beginning of disease management.

**Fig. S8**. Dynamics of observations, tree and orchard removals, under different management strategies (French management in orchards, or economically improved strategies) and epidemic contexts (reference context, or harsher epidemics: half the duration of the expected latent period, *θ_exp_*, or doubled transmission coefficient, *β*) in 100 simulations. The dashed vertical line indicates the beginning of disease management.

**Fig. S9**. Boxplots of *Y* (equivalent number of fully productive trees per hectare and per year in A and C), and *NPV* (net present value in B and D) after 30 years of management in harsher epidemic contexts (half the duration of the expected latent period, *θ_exp_*, in A and B; or doubled transmission coefficient, *β*, in C and D). Different scenarios are simulated: absence of disease, absence of management, disease managed with the reference strategy (French management in orchards), or with economically improved management strategies identified through two different methods (combination associated with the best value or the highest percentile of *μNPV*).

**Fig. S10**. Boxplots of *Y* (equivalent number of fully productive trees per hectare and per year in A and C), and *NPV* (net present value in B and D) after 30 years of management in harsh epidemic contexts (half the duration of the expected latent period, *θ_exp_*, in A and B; or doubled transmission coefficient, *β*, in C and D), with strategies specifically improved in these contexts. Different scenarios are simulated: absence of disease, absence of management, disease managed with the reference strategy (French management in orchards), or with economically improved management strategies identified through two different methods (combination associated with the best value or the highest percentile of *μNPV*).

**Fig. S11**. Dynamics of annual prevalence (blue curves) and incidence (red curves) under management strategies specifically improved in harsh epidemic contexts (half the duration of the expected latent period, *θ_exp_*, or doubled coefficient of transmission, *β*) in 100 simulations. The dashed vertical line indicates the beginning of disease management.

**TABLE S1**. Economic analysis of prunus cultivation in France. This analysis enables to estimate economic parameters (yellow cells: *p, c_H_, c_S_, c_R_, c_F_, X_SEHD_*) used to compute the economic criterion *(NPV)*, from parameters informed by available data (fruit sale price, grey cell) or expert opinion (green cells). The analysis is based on the cultivation of an average 1-ha orchard during 15 years in the absence of PPV and without discounting (i.e., equivalent to the cultivation of 15 asynchronous orchards during one year). White cells are computed from other cells.

