## Supporting Information for "Improving management strategies of plant diseases using sequential sensitivity analyses"

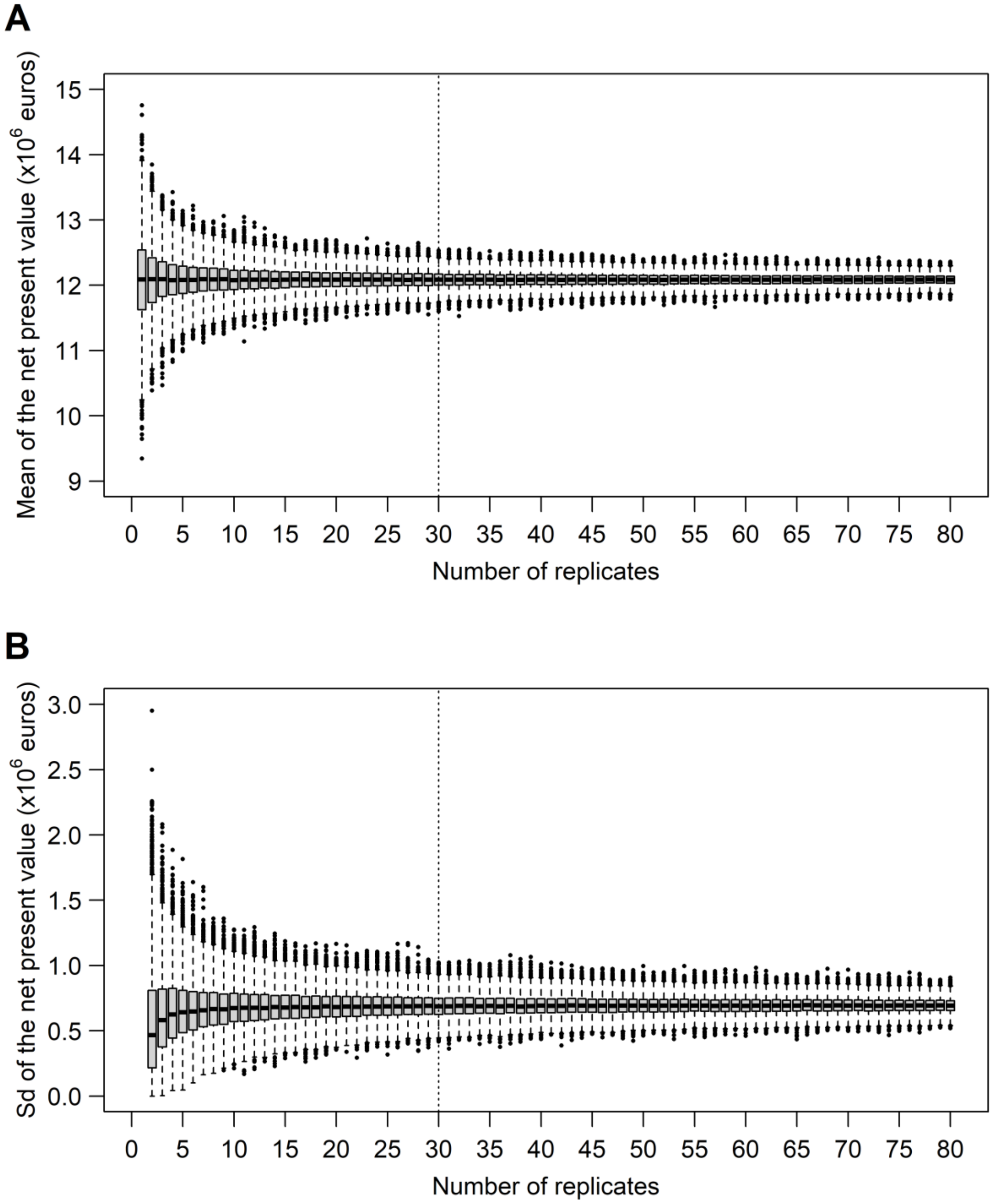


**Fig. S1.** Distribution of 5,000 means (A) and standard deviations (B) of the model output (net present value, $NPV$) computed with an increasing number of replicates. All model parameters are fixed at their reference value. The dashed vertical line delimits the selected number of replicates (30).

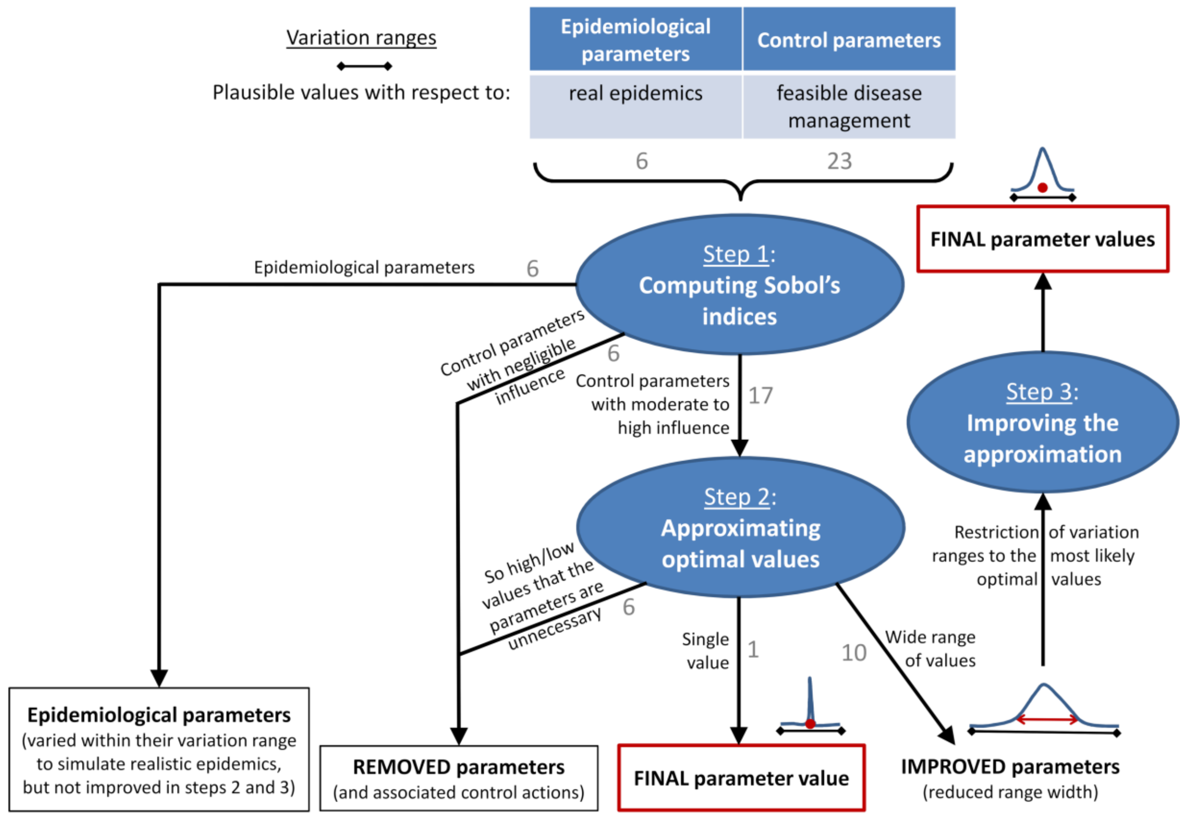


**Fig. S2.** Schematic representation of the heuristic approach to improve sharka management using sequential sensitivity analyses of a simulation model. In every sensitivity analysis (SA), variation ranges of epidemiological parameters are restricted to simulate realistic epidemics, whereas control parameters are varied within values that are compatible with a feasible management. In step 1, a first SA highlights key epidemiological and control parameters. In step 2, a second SA targets the most influential control parameters and approximates optimal values. Then, parameters associated with a wide range of improved values are further improved via a dedicated SA in step 3. Control actions governed by parameters with negligible influence (in step 1) or associated with unrealistic values (in step 2) can be removed from the model. Blue ellipses: sensitivity analyses; grey numbers: number of parameters in the model; boxes: outcomes of the approach.

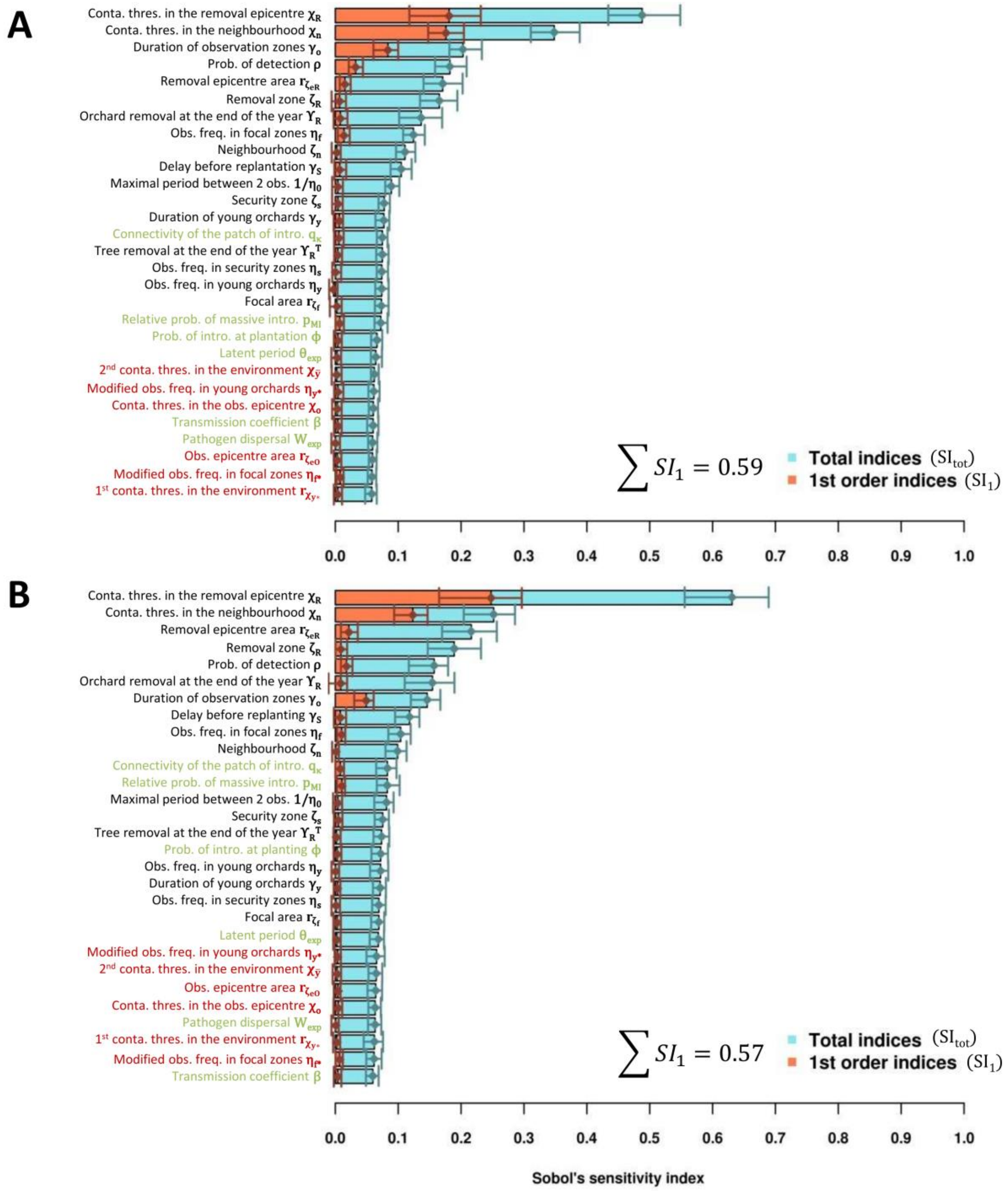


**Fig. S3.** Step 1: Sobol's sensitivity indices of the 23 control parameters and 6 epidemiological parameters on the standard deviation of the stochastic replicates (A: $\sigma_Y$, average number of fully productive trees; B: $\sigma_{NPV}$, average net present value). Parameters in black are kept in step 2; parameters in red are removed; parameters in green are the epidemiological parameters.

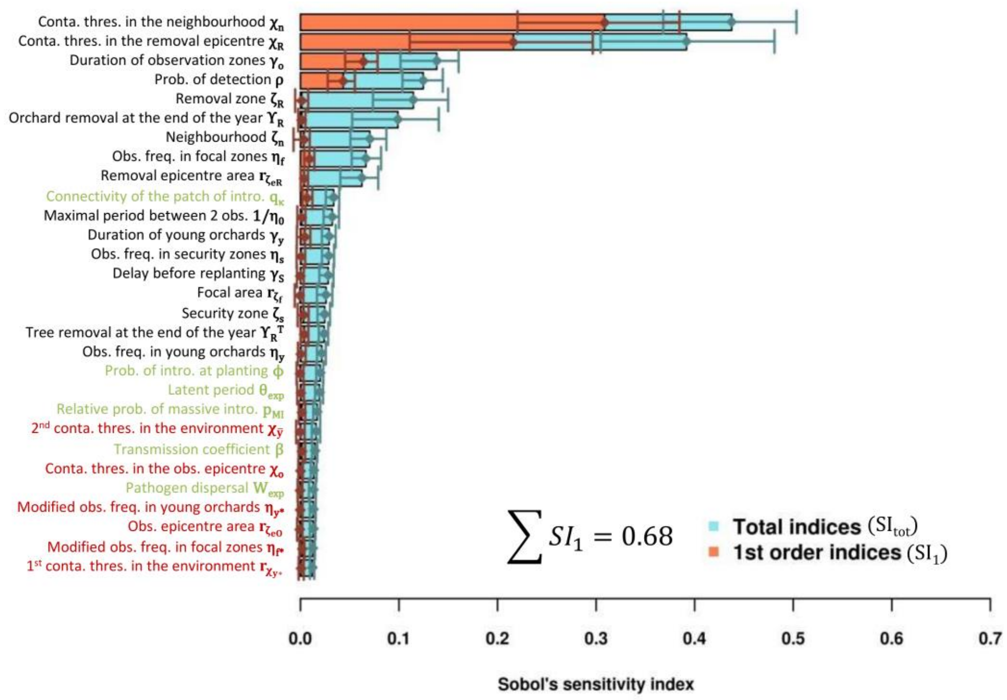


**Fig. S4.** Step 1: Sobol's sensitivity indices of the 23 control parameters and 6 epidemiological parameters on the mean epidemiological output of the stochastic replicates ($\mu_Y$, average number of fully productive trees). Parameters in black are kept in step 2; parameters in red are removed; parameters in green are the epidemiological parameters. Results on the economic output (average net present value, $\mu_{NPV}$) are presented in Fig. 2.

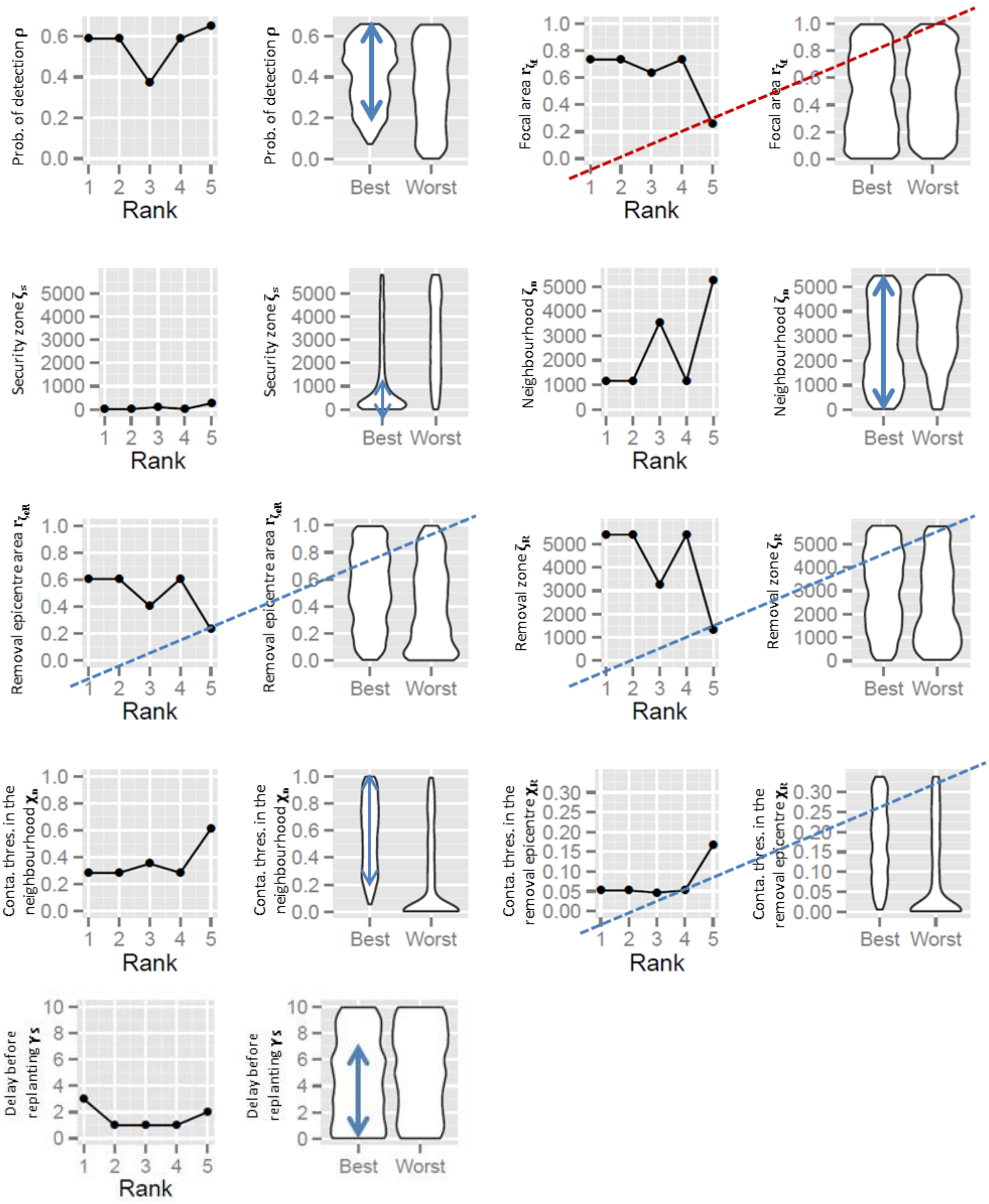


(legend on next page)

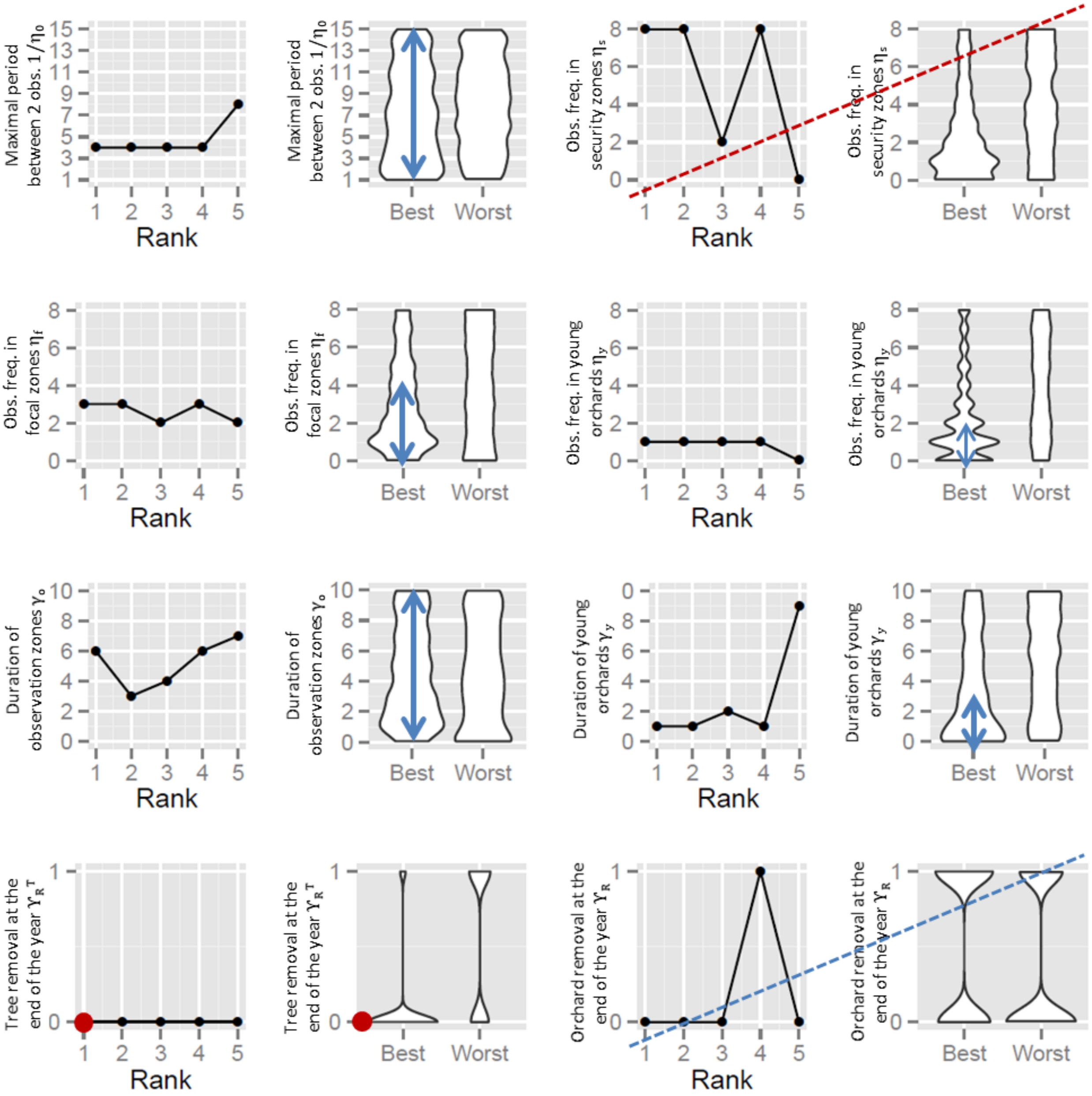


**Fig. S5.** Step 2: Five best values (first and third columns), and distribution of each parameter for the highest and lowest percentiles (second and last columns) of the economic criterion ($\mu_{NPV}$, average net present value) in the second sensitivity analysis. A single improved value is found for 1 parameter (red circle), 6 unnecessary parameters can be removed (dashed red lines: the focal zone matches the security zone, so both zones can be merged; dashed blue lines: improved values for removal parameters are so high that removals might never occur) and the 10 remaining parameters are to be further improved in step 3, using restricted variation ranges (blue arrows).

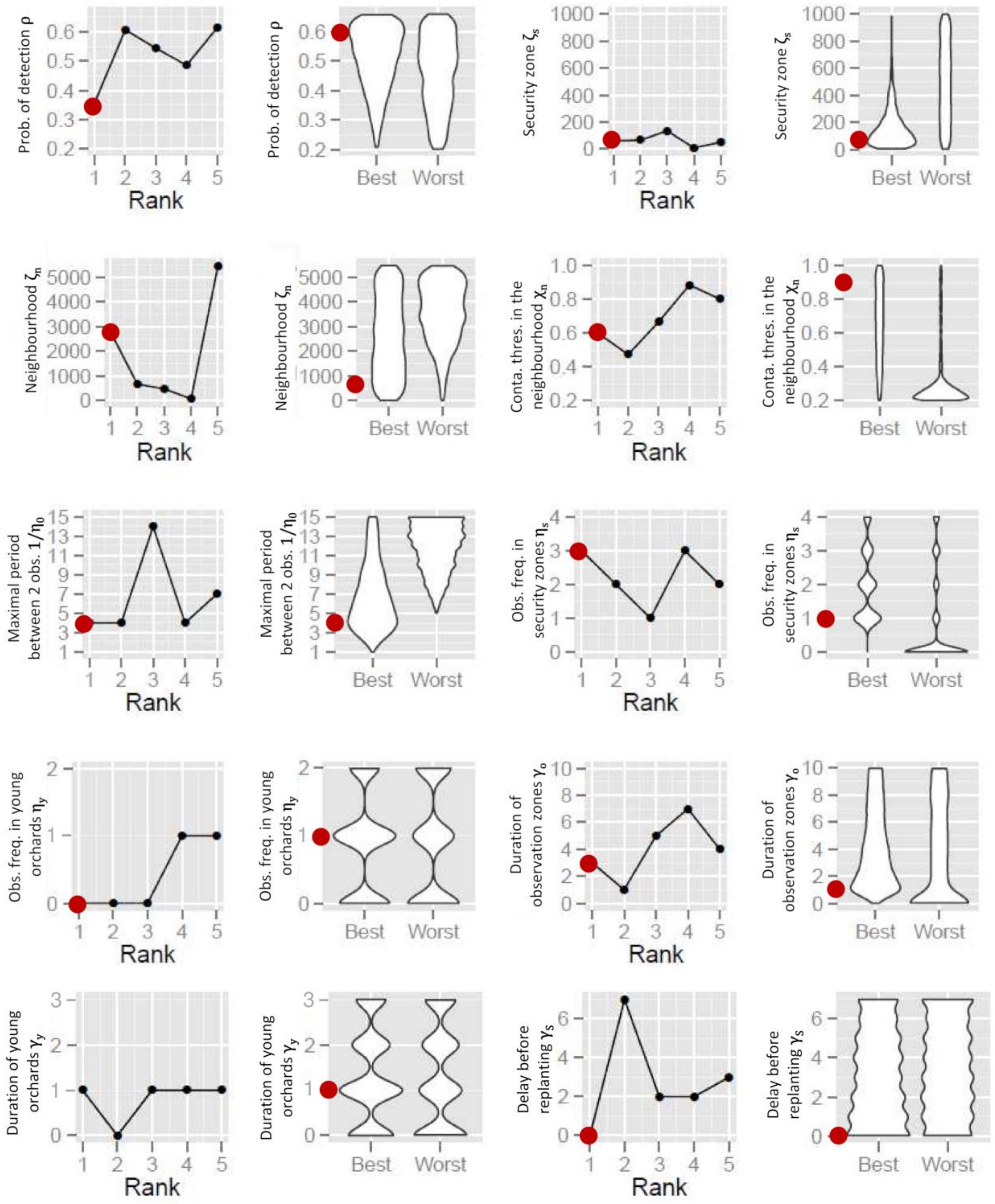


**Fig. S6.** Step 3: Five best values (first and third columns), and distribution of each parameter for the highest and lowest percentiles (second and last columns) of the economic criterion ($\mu_{NPV}$, average net present value) in the third sensitivity analysis. Improved values for each control parameter are indicated by red dots. The duration of young orchards ($\gamma_y$) is irrelevant in the best-value strategy because the observation frequency in young orchards ($\eta_y$) is 0.

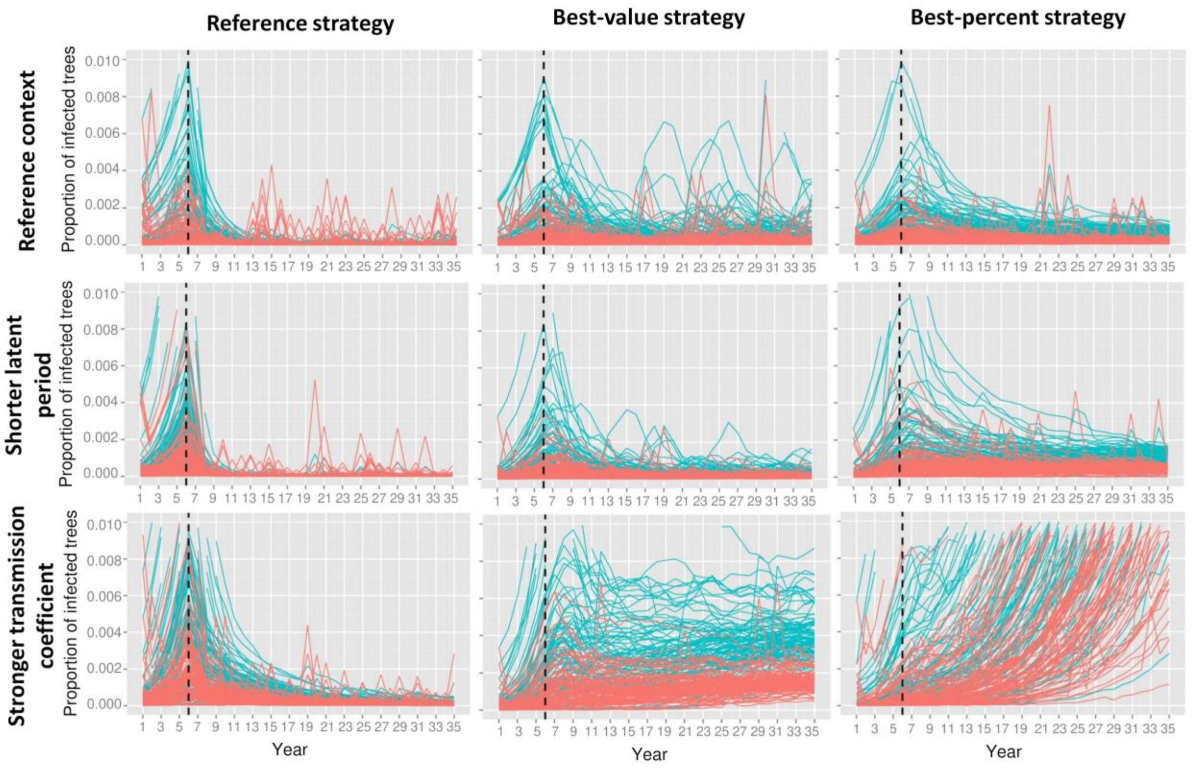


**Fig. S7.** Dynamics of annual prevalence (blue curves) and incidence (red curves) under different management strategies (French management in orchards, or economically improved strategies) and epidemic contexts (reference context, or harsher epidemics: half the duration of the expected latent period, $\theta_{exp}$, or doubled transmission coefficient, $\beta$) in 100 simulations. The dashed vertical line indicates the beginning of disease management.

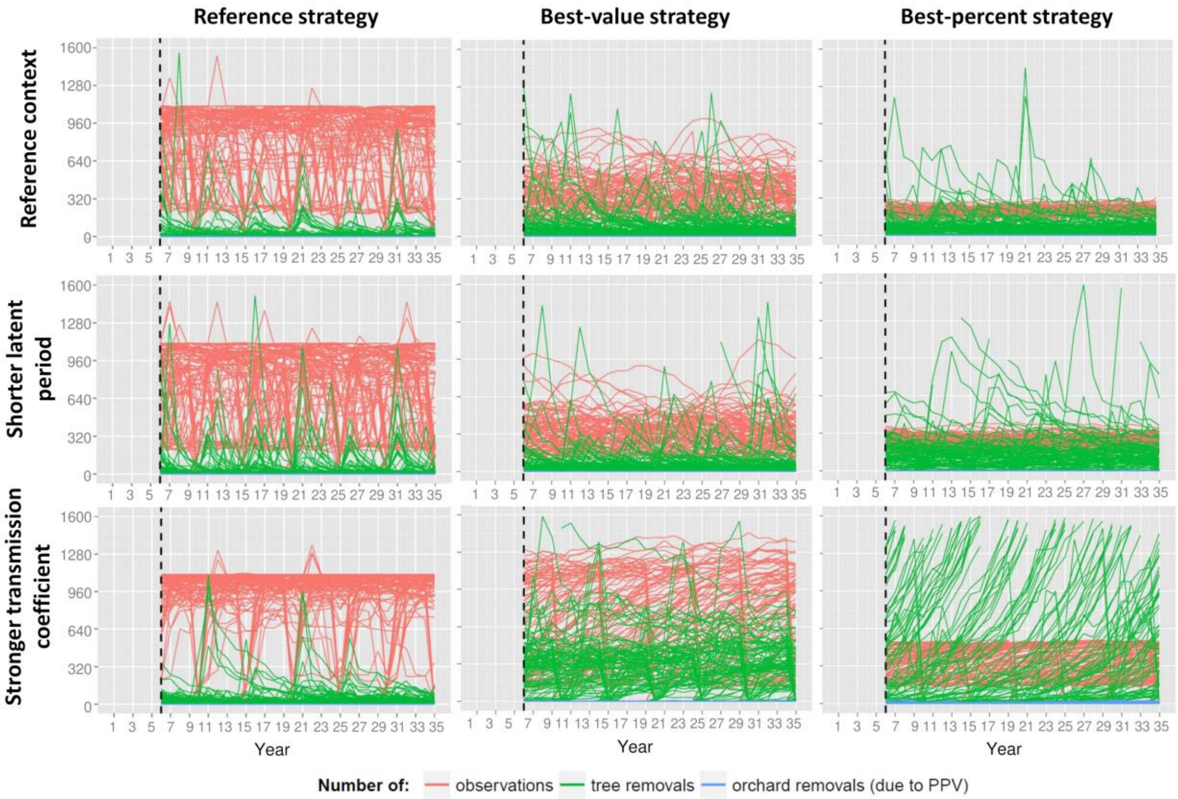


**Fig. S8.** Dynamics of observations, tree and orchard removals, under different management strategies (French management in orchards, or economically improved strategies) and epidemic contexts (reference context, or harsher epidemics: half the duration of the expected latent period, $\theta_{exp}$, or doubled transmission coefficient, $\beta$) in 100 simulations. The dashed vertical line indicates the beginning of disease management.

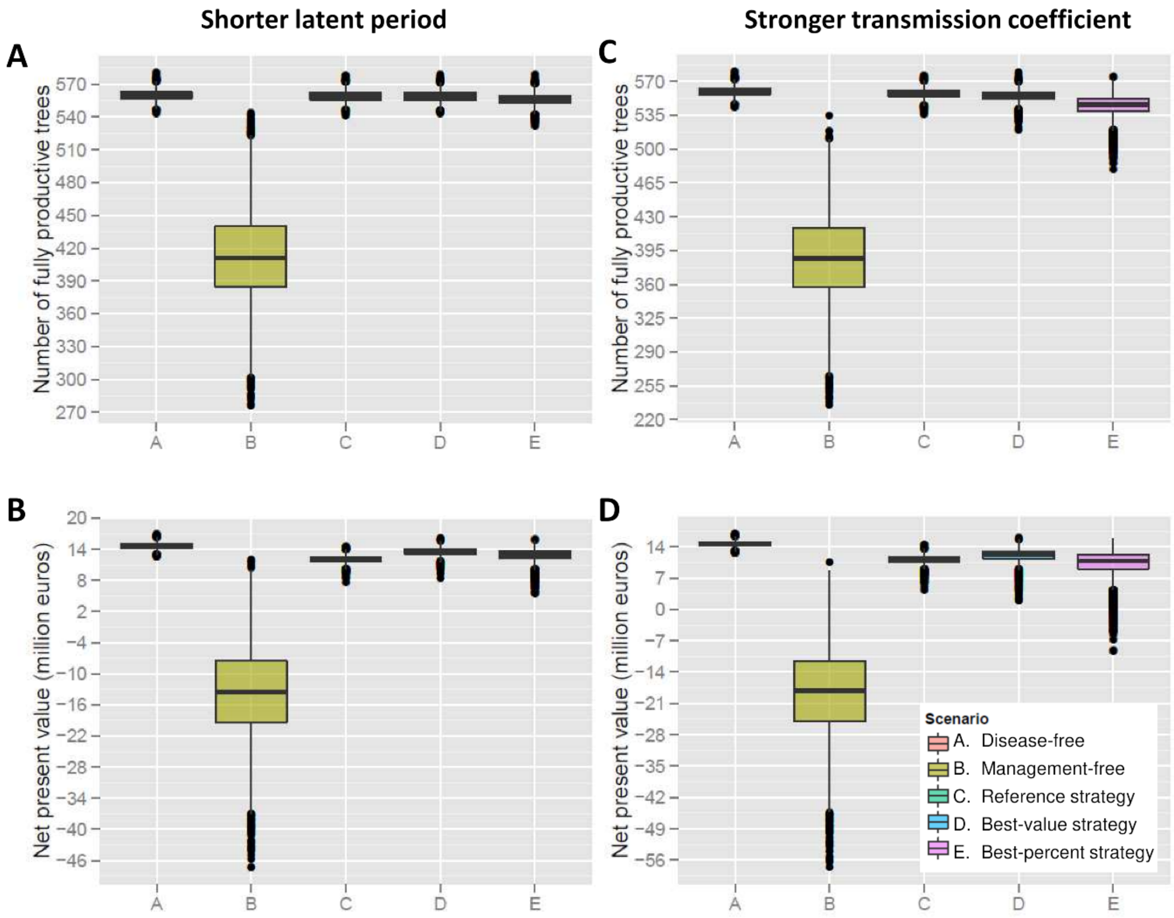


**Fig. S9.** Boxplots of $Y$ (equivalent number of fully productive trees per hectare and per year in A and C), and $NPV$ (net present value in B and D) after 30 years of management in harsher epidemic contexts (half the duration of the expected latent period, $\theta_{exp}$, in A and B; or doubled transmission coefficient, $\beta$, in C and D). Different scenarios are simulated: absence of disease, absence of management, disease managed with the reference strategy (French management in orchards), or with economically improved management strategies identified through two different methods (combination associated with the best value or the highest percentile of $\mu_{NPV}$).

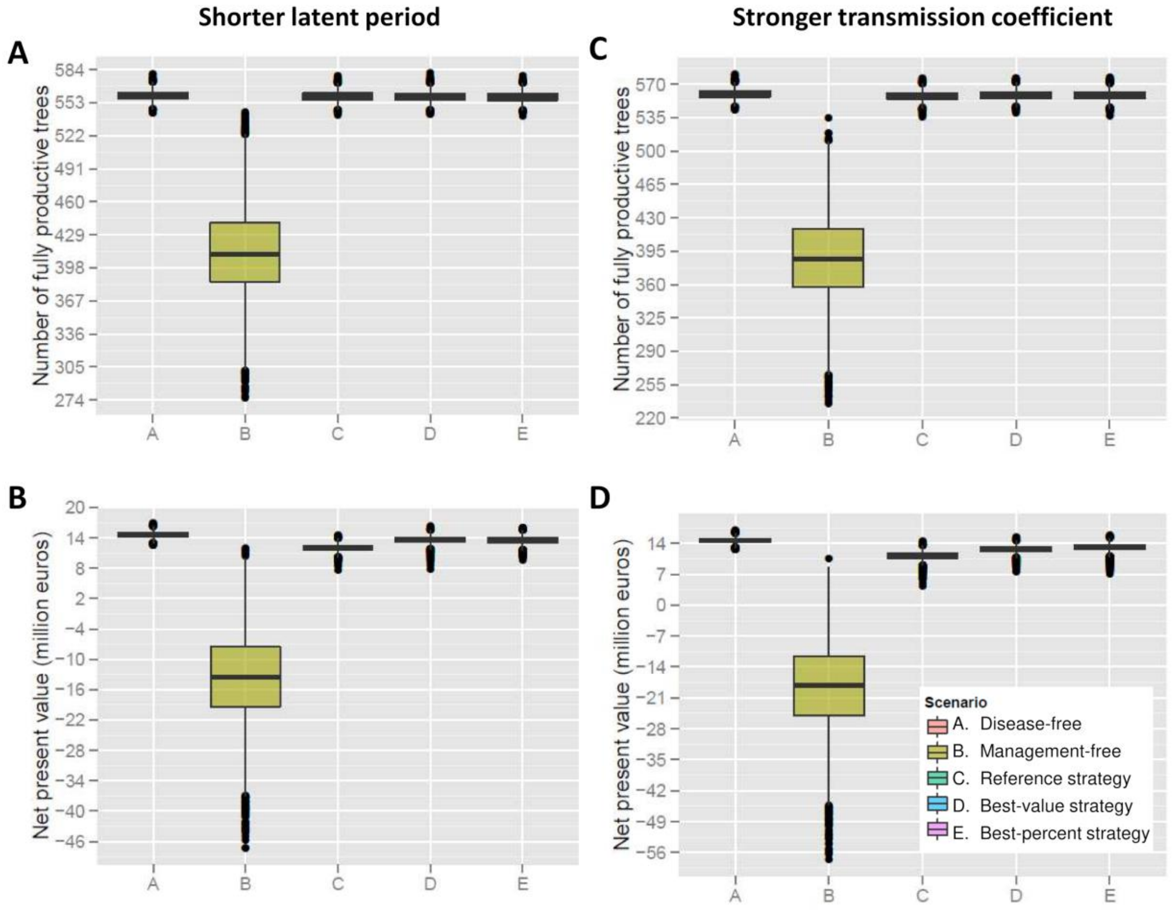


**Fig. S10.** Boxplots of $Y$ (equivalent number of fully productive trees per hectare and per year in A and C), and $NPV$ (net present value in B and D) after 30 years of management in harsh epidemic contexts (half the duration of the expected latent period, $\theta_{exp}$, in A and B; or doubled transmission coefficient, $\beta$, in C and D), with strategies specifically improved in these contexts. Different scenarios are simulated: absence of disease, absence of management, disease managed with the reference strategy (French management in orchards), or with economically improved management strategies identified through two different methods (combination associated with the best value or the highest percentile of $\mu_{NPV}$).

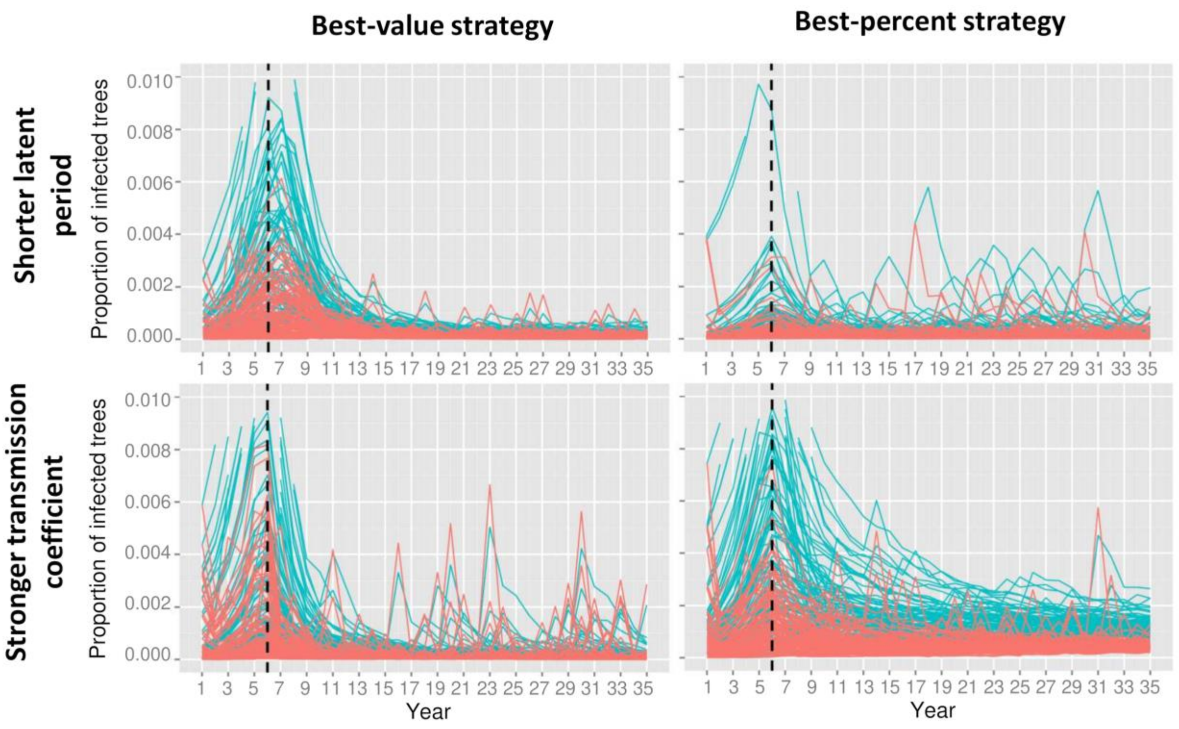


**Fig. S11.** Dynamics of annual prevalence (blue curves) and incidence (red curves) under management strategies specifically improved in harsh epidemic contexts (half the duration of the expected latent period, $\theta_{exp}$, or doubled coefficient of transmission, $\beta$) in 100 simulations. The dashed vertical line indicates the beginning of disease management.

| Expert opinion | Parameter | Value | References and comments |
|---|---|---|---|
| Available data | Fruit sale price (€/kg): | 1,49 € | Agreste 2013, 2014, 2015; FranceAgriMer 2015; INSEE 2015 |
| Parameter of interest | Gross margin of sold fruits (€/kg): | 0,10 € | (includes direct and fixed costs like amortization of the orchard) |
| | Proportion of unsold fruits: | 17% | |
| | Cost price of produced fruits (€/kg): | 1,16 € | |
| | Total expenses of an orchard (€/ha): | 417 000 € | |
| | Proportion harvest/packaging in total expenses: | 47% | (0.15 €/kg for harvest, 0.30 €/kg for wrapping materials, 0.10 €/kg for packaging) |
| | Total expenses without harvests (€/ha): | 219 000 € | |
| | Total expenses without planting, harvests and removal (€/ha): | 204 000 € | |
| | Profitability threshold ($\chi_{SEHD}$): | 66% | |

| | | Maximal fruit yield: | Maximal fruit sales: | Maximal harvest expenses: | Planting cost: | Removal cost: | Yearly expenses without planting, harvests and removal: | | Average gross margin |
|---|---|---|---|---|---|---|---|---|---|
| | | (kg/ha/an) | p (€/ha) | $c_H$ (€/ha/harvest) | $c_S$ (€/ha) | $c_R$ (€/ha) | $c_F$ (€/ha) | | (€/ha/an) |
| | | 25 000 | 37 250 € | 16 500 € | 14 000 € | 1 000 € | 13 600 € | | 2000 |
| **Orchard age (years)** | **Relative fruit yield: *y*** | **Fruit production (kg/ha)** | **Fruit sales (€/ha)** | **Harvest expenses (€/ha)** | **Planting expenses (€/ha)** | **Removal expenses (€/ha)** | **Fixed expenses (€/ha)** | **Total expenses without PPV (€/ha)** | **Gross margin without PPV (€/ha)** |
| 1 | 0% | 0 | 0 € | 0 € | 14 000 € | 0 € | 13 600 € | 27 600 € | -27 600 € |
| 2 | 0% | 0 | 0 € | 0 € | 0 € | 0 € | 13 600 € | 13 600 € | -13 600 € |
| 3 | 50% | 12 500 | 18 625 € | 8 250 € | 0 € | 0 € | 13 600 € | 21 850 € | -3 225 € |
| 4 | 65% | 16 250 | 24 213 € | 10 725 € | 0 € | 0 € | 13 600 € | 24 325 € | -112 € |
| 5 | 85% | 21 250 | 31 663 € | 14 025 € | 0 € | 0 € | 13 600 € | 27 625 € | 4 038 € |
| 6 | 100% | 25 000 | 37 250 € | 16 500 € | 0 € | 0 € | 13 600 € | 30 100 € | 7 150 € |
| 7 | 100% | 25 000 | 37 250 € | 16 500 € | 0 € | 0 € | 13 600 € | 30 100 € | 7 150 € |
| 8 | 100% | 25 000 | 37 250 € | 16 500 € | 0 € | 0 € | 13 600 € | 30 100 € | 7 150 € |
| 9 | 100% | 25 000 | 37 250 € | 16 500 € | 0 € | 0 € | 13 600 € | 30 100 € | 7 150 € |
| 10 | 100% | 25 000 | 37 250 € | 16 500 € | 0 € | 0 € | 13 600 € | 30 100 € | 7 150 € |
| 11 | 100% | 25 000 | 37 250 € | 16 500 € | 0 € | 0 € | 13 600 € | 30 100 € | 7 150 € |
| 12 | 100% | 25 000 | 37 250 € | 16 500 € | 0 € | 0 € | 13 600 € | 30 100 € | 7 150 € |
| 13 | 100% | 25 000 | 37 250 € | 16 500 € | 0 € | 0 € | 13 600 € | 30 100 € | 7 150 € |
| 14 | 100% | 25 000 | 37 250 € | 16 500 € | 0 € | 0 € | 13 600 € | 30 100 € | 7 150 € |
| 15 | 100% | 25 000 | 37 250 € | 16 500 € | 0 € | 1 000 € | 13 600 € | 31 100 € | 6 150 € |
| **15** | **12** | **300 000** | **447 000 €** | **198 000 €** | **14 000 €** | **1 000 €** | **204 000 €** | **417 000 €** | **30 000 €** |

**TABLE S1.** Economic analysis of prunus cultivation in France. This analysis enables to estimate economic parameters (yellow cells: $p$, $c_H$, $c_S$, $c_R$, $c_F$, $X_{SEHD}$) used to compute the economic criterion ($NPV$), from parameters informed by available data (fruit sale price, grey cell) or expert opinion (green cells). The analysis is based on the cultivation of an average 1-ha orchard during 15 years in the absence of PPV and without discounting (i.e., equivalent to the cultivation of 15 asynchronous orchards during one year). White cells are computed from other cells.
